# PYK2 controls intestinal inflammation via activation of IRF5 in macrophages

**DOI:** 10.1101/2020.05.24.113076

**Authors:** Grigory Ryzhakov, Hannah Almuttaqi, Alastair L. Corbin, Tariq Khoyratty, Dorothee Berthold, Samuel Bullers, Hayley L Eames, Zhichao Ai, Sarah Bonham, Roman Fischer, Luke Jostins-Dean, Simon P.L. Travis, Benedikt M. Kessler, Irina A. Udalova

## Abstract

Inflammatory bowel disease (IBD) is a group of inflammatory disorders of the gastro-intestinal tract caused by a complex combination of genetic and environmental factors. Interferon regulating factor 5 (IRF5) is a multifunctional regulator of immune responses, which plays a key pathogenic role in mouse colitis models and is a genetic risk factor for IBD. A screen of a protein kinase inhibitor library in macrophages revealed a list of putative IRF5 kinases. Among the top hits validated in multiple *in vitro* assays, protein-tyrosine kinase 2-beta (PTK2B or PYK2) was identified as the only IBD genetic risk factor, known to impact gene expression in myeloid cells^1,2^. Phospho-proteomics and mutagenesis analyses established that PYK2 directly phosphorylates and activates IRF5 at tyrosine (Y) 171. IRF5 nuclear translocation and recruitment to target genes was impaired in PYK2-deficient cells or in cells treated with PYK2 inhibitors. Importantly, macrophage transcriptomic signature under PYK2 inhibition phenocopied IRF5 deficiency. Treatment with a PYK2 inhibitor reduced pathology and inflammatory cytokine production in *Helicobacter hepaticus* + anti-IL-10R antibody induced colitis model. It also decreased levels of pro-inflammatory cytokines in human colon biopsies taken from patients with ulcerative colitis. Thus, we have identified a major role for PYK2 in regulating the inflammatory response and mapped its activity to the IRF5 innate sensing pathway, opening opportunities for therapeutic interference with it in IBD and other inflammatory conditions.

---

A recent single-cell transcriptomic analysis of colon biopsies from patients with ulcerative colitis (UC) provided a framework for linking GWAS risk loci with specific cell types and functional pathways and helped to nominate causal genes across GWAS loci^3^, amongst them Interferon regulatory factor 5 (IRF5). IRF5 is a multifunctional regulator of immune responses^4–6^. The IRF5 risk variant has consistent effects across monocytes and macrophage conditions, but also impacts gene expression and splicing across a wide range of other immune cells and tissues^7^.

Recent studies using IRF5-deficient mice have established a critical role of this transcription factor in the pathogenesis of mouse models of colitis^8,9^. IRF5 is proposed to exert its molecular function via a cascade of events involving its phosphorylation, ubiquitination, dimerisation, nuclear translocation and selective binding to its target genes to enable their expression^10^. Despite its known physiological role, the molecular mechanisms of IRF5 activation are still debated. Several kinases including TBK1, RIP2, IKKε, IRAK4, TAK1, and IKKβ have been proposed to phosphorylate and activate IRF5^11–17^, while IKKα inhibits IRF5^18^. Lyn, a Src family kinase has been shown negatively to regulate IRF5 in the TLR-MyD88 pathway in a kinase independent manner via direct binding to IRF5^19^.

In this study, we identified another nominated causal gene for UC^3^, Protein Tyrosine Kinase 2b (PTK2B/PYK2), as a key regulator of IRF5 activation, macrophage inflammatory response, and intestinal pathology extending its currently accepted function in macrophage morphology and migration^20^.

We have previously established an *in vitro* reporter system for measuring IRF5-dependent transcription based on the TNF (IRF5-dependent gene)-promoter driven luciferase construct, which contains a number of interferon-stimulated response elements (ISREs)^21^. In our hands the TNF promoter reporter consistently showed a stronger response to IRF5 than the standardly used ISRE-luciferase reporter (**Supplementary Fig. 1a, b**), either due to the number of ISRE sites and/or previously reported IRF5 cooperation with NF-kB RelA^22^. This reporter system was used to screen a library of small molecules^23^, for which inhibitory properties against 221 protein kinases in the Protein Kinase Inhibitors Screen (PKIS) have been established (**Supplementary Fig. 1c, d, e**). After the first screen in RAW264.7 macrophages and 2 rounds of re-screening using different cell types and three different inhibitor concentrations, we composed the final list of 34 candidate IRF5 kinases, among which TBK1, IKKe and IRAK4 were previously proposed to target IRF5^12,16,24^ (**Fig. 1a, Source Data 1)**.

**Figure 1.**
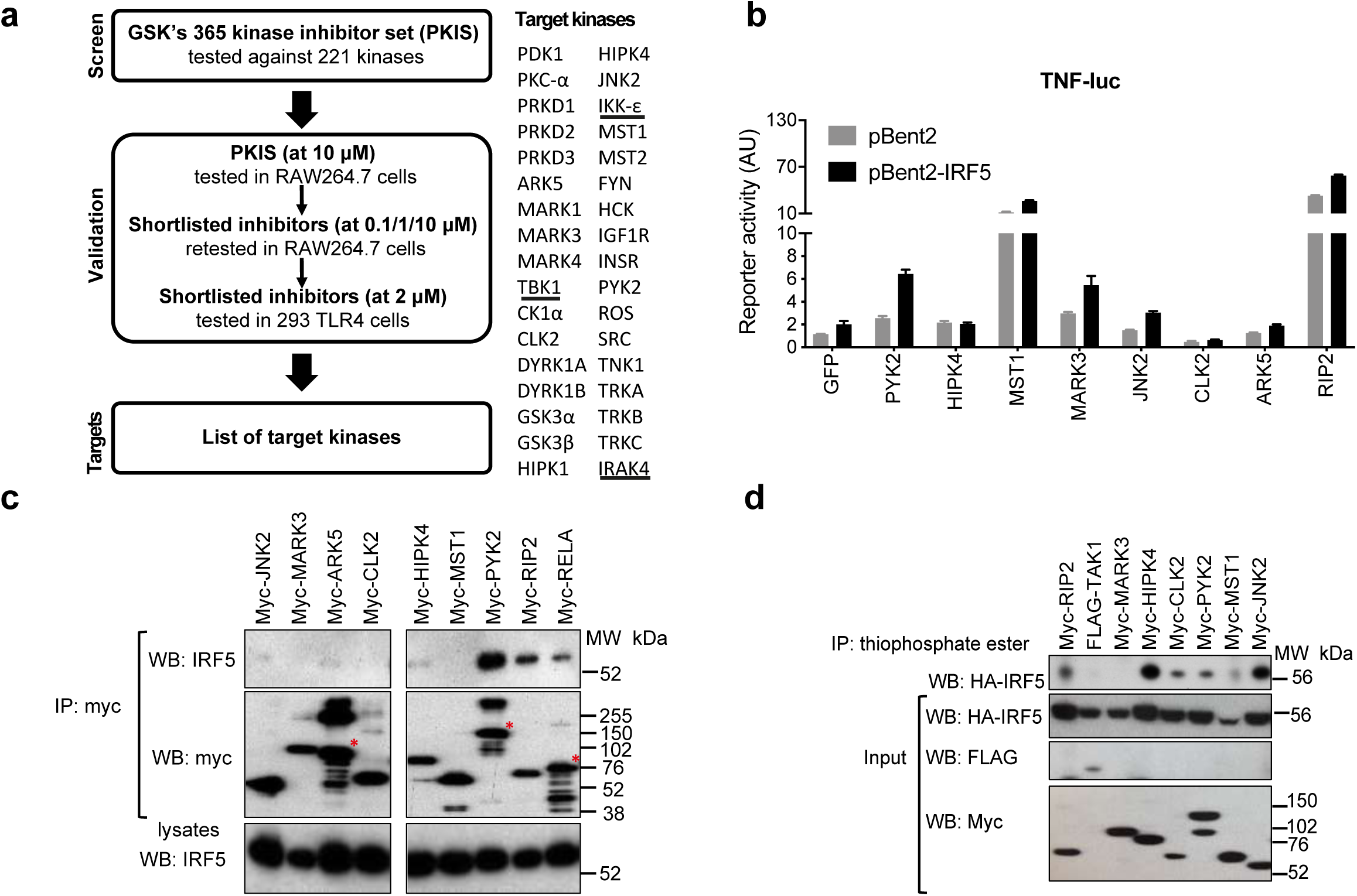
Small molecule library screening and in vitro validation of shortlisted candidate IRF5 kinases confirms Pyk2 as a positive hit. (**a**) The screening workflow showing the initial large screening in RAW264.7 cells, and the subsequent screens in RAW264.7 and 293 TLR4 cells. The top inhibitors were shortlisted based on their efficacy towards IRF5 reporter and low toxicity. Based on the known activities of these molecules against 221 kinases in the PKIS set, 34 kinases affected by the top inhibitors were shortlisted. The underlined kinases were previously proposed to target IRF5. **(b)** Impact of the kinases on the IRF5 reporter activity. Luciferase activities were measured in cells co-expressing IRF5 (or empty plasmid control, pBent2), TNF-luc reporter and one of the shortlisted kinases. Reporter activity was calculated as firefly luciferase activity normalised against constitutively expressed Renilla luciferase units and is shown as compared to the values in cells not expressing any kinase. **(c)** Binding of IRF5 to the shortlisted kinases. Myc-tagged kinases and HA-tagged IRF5 were co-expressed in 293 ET cells. Cell lysates were subjected to immunoprecipitation (IP) using anti-myc antibody and levels of kinases and IRF5 in the IP eluates and proteins were determined by Western blot. Asterisks indicates expected molecular weight. (**d**) *In vitro* kinase assays of 293 ET cells co-transfected with HA-IRF5 and myc- or flag-tagged kinases. Proteins in the pull-downs and lysates were detected by Western blotting using antibodies against HA- (IRF5) and myc- and FLAG- (kinases).

For further functional validation we selected poorly explored proteins or those with known links to inflammatory processes – PYK2, HIPK4, ARK5, CLK2, MARK3, JNK2 and MST1. We then cloned the cDNAs encoding them into mammalian expression vectors and tested them in functional assays. As controls, we included into these assays RIP2 kinase, the known intermediate in the IRF5-dependent innate immune signalling pathways^11^. When we overexpressed the kinases with IRF5 and TNF-luciferase reporter in HEK-293 TLR4/CD14/MD-2 cells, we found that overexpression of PYK2, JNK2 or MARK3, boosted IRF5-dependent TNF-reporter activation (**Fig. 1b**). Similar to the known IRF5 binding partners RIP2 and RelA^11,22,25^, PYK2 could strongly bind IRF5 in co-immunoprecipitation assays, while HIPK4, ARK5 and JNK2 showed only weak association with IRF5 (**Fig. 1c**). We further tested the ability of these kinases to phosphorylate overexpressed IRF5 in 293 ET cell lysates (**Supplementary Fig. 1f**). We were able to detect phosphorylated IRF5 in the presence of HIPK4, CLK2, JNK2, MST1, PYK2 and RIP2 as a positive control (**Fig. 1d**). Lastly, we examined the evidence of genetic association of the selected kinases with IBD and found that PYK2 was the only known genetic risk factor^26^. Moreover, the risk variant for PYK2 was shown to impact gene expression in monocytes and macrophages^27^. Based on observed functional interactions with IRF5 and genetic association with IBD, PYK2 was singled out for further investigation in macrophages (**Supplementary Fig. 1g**).

In RAW264.7 macrophages we could detect PYK2 binding to IRF5 at the endogenous level (**Fig. 2a**). In line with previous studies, we also found LPS-induced PYK2 phosphorylation on Y402 (**Fig. 2b**)^28,29^. To investigate the kinase’s role in the TLR4/IRF5 signalling axis, we generated IRF5- and PYK2-deficient mouse RAW264.7 macrophages using a CRISPR-Cas9 approach (**Supplementary Fig. 2a**). First, we explored the impact of PYK2 deficiency on IRF5-dependent signalling by transfecting wild type (WT) and PYK2-deficient RAW264.7 macrophages with IRF5-expressing and the TNF-promoter driven luciferase plasmid. We observed a marked reduction of the LPS-induced reporter activity in IRF5 expressing cells lacking PYK2 (**Fig. 2c**). When we expressed recombinant PYK2 in PYK2-deficient cells, we achieved a partial reconstitution of the PYK2 levels in RAW264.7 macrophages and a partial restoration of the reporter activity (**Supplementary Fig. 2b, c**). Next, we examined if PYK2 deficiency would directly impact IRF5 activation and function by measuring IRF5 recruitment to its target gene promoter and enhancer regions using chromatin immunoprecipitation (ChIP) assay^4,21^. IRF5 recruitment to *Il6, Il1a* and *Tnf* gene promoters was impaired in PYK2 knockout cells (**Fig. 2d, Supplementary Fig. 2d**). Consequently, we observed attenuated recruitment of RNA polymerase II at the same promoters indicating reduced gene transcription (**Fig. 2d, Supplementary Fig. 2d**). We also detected reduction in mRNA induction of these cytokines, as well as chemokines *Ccl4, Ccl5*, by LPS in PYK2-deficient cells, comparable to or even stronger than in IRF5 knockout cells (**Fig. 2e, Supplementary Fig. 2e**). Conversely, LPS-induced IL-10 induction was increased in PYK2 knockout (**Supplementary Fig. 2e**), similarly to our previous findings in IRF5 knockout cells ^4^.

**Figure 2.**
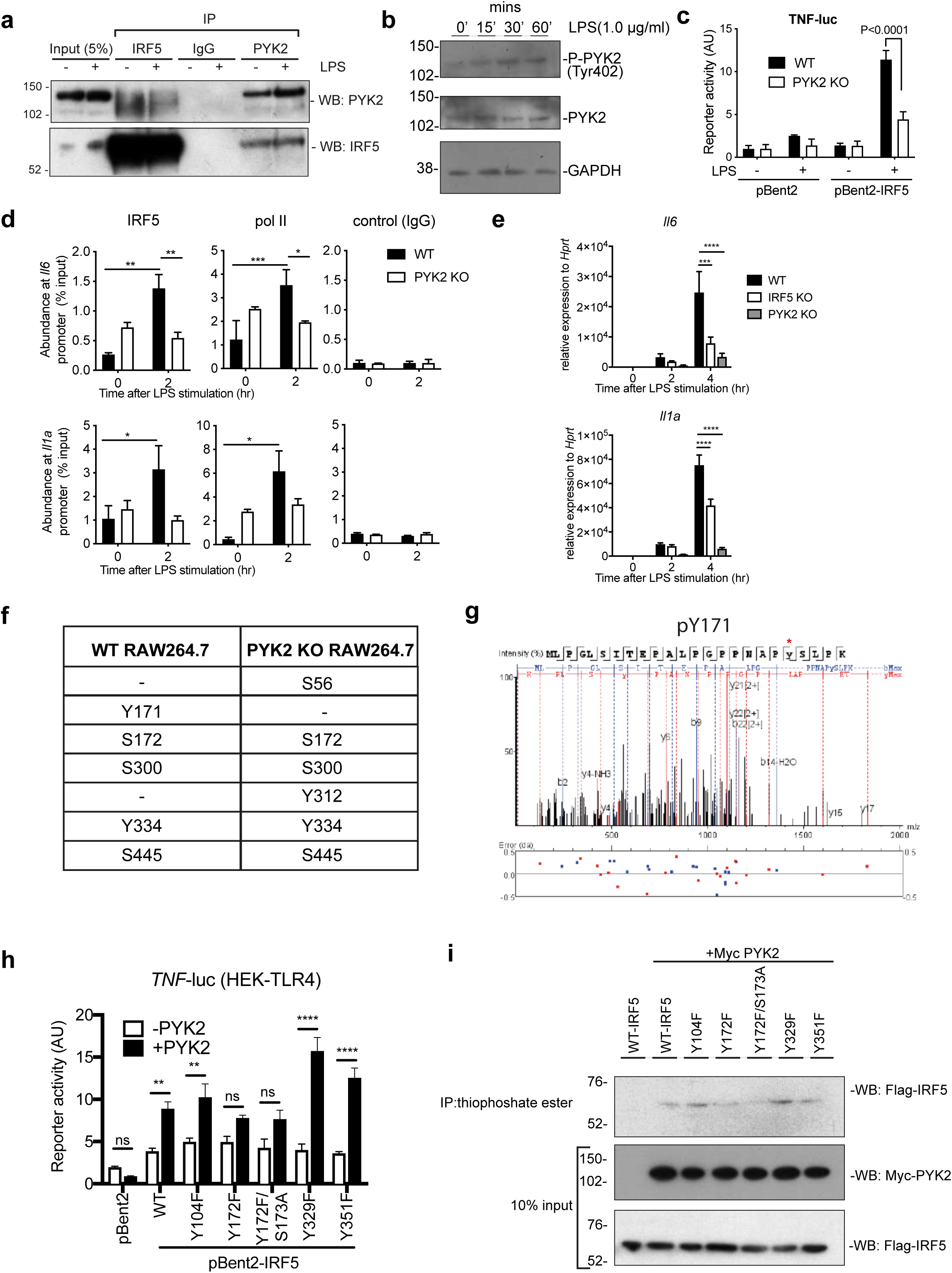
PYK2 regulates IRF5 activation and IRF5-mediated transcription. (**a**) Endogenous co-immunoprecipitation in RAW264.7 macrophages. Cells were stimulated with LPS for 10 minutes and immunoprecipitated with IRF5, PYK2 or an isotype control antibody. Immunoprecipitates were eluted from IP beads and proteins present in cell lysates (5% inputs) and eluates were detected by immunoblotting with antibodies against IRF5 or PYK2. (**b**) Immunoblot of LPS-induced PYK2 tyrosine phosphorylation. Blots were probed with antibodies specific for PYK2 phosphorylated on Tyr-402, total PYK2 and GAPDH. (**c**) TNF-luc reporter activity in the absence or presence of ectopically expressed IRF5 in wild type and PYK2 KO RAW264.7 cells stimulated with LPS 6 hrs or left untreated. (**d**) IRF5 and pol II binding to *Il6* and *Il1a* gene promoter in resting or LPS-treated (2h, 500 ng/ml) wild type or PYK2 KO RAW264.7 cells as measured by the chromatin immunoprecipitation (ChIP) method. A non-specific IgG antibody was used as a negative control for ChIP. (**e**) *Il6* and *Il1a* mRNA induction in wild type, PYK2 KO or IRF5 KO RAW264.7 cells stimulated with LPS (500 ng/ml) for 0, 2, or 4 hrs. Gene expression was measured by qPCR. **(f)** Phosphorylation sites identified in LPS-stimulated WT and PYK2 KO RAW264.7 cells. **(g)** MS/MS spectrum of the IRF5 derived tryptic peptide 152-179 indicating phosphorylation at positions Y171. Fragmentation ions of the b- and y-series are indicated in blue and red, respectively. **(h)** Phospho-site inactivating mutations were introduced in IRF5 (human, isoform v2) in HEK-TLR4 cells and their effect on the TNF-luciferase reporter assay was measured in the absence or presence of PYK2. Reporter activity is expressed as firefly luciferase levels relative to Renilla levels and values are means of three independent experiments. **(i)** *In vitro* kinase assay and immunoblot of IRF5-site specific tyrosine mutants. HEKTLR4 cells were co-transfected with FLAG-IRF5 tyrosine mutants as indicated and Myc-PYK2. 10% lysate was kept for input and the remaining used for *in vitro* kinase reactions. Kinase assays were detected by western blot using antibodies against Flag-(IRF5) and Myc-(PYK2). All values in (c-e, h) are shown as mean values +/− SEM from n=3 experiments. Comparison by two-way ANOVA \**P*<0.05, \*\**P*<0.01, \*\*\**P*<0.001, and \*\*\*\**P*<0.0001.

To validate our observations in primary cells, we utilised immortalised myeloid progenitor HoxB8 cells^30^, which differentiated into non-proliferating mature macrophages after GM-CSF-induced differentiation for 5 days (**Supplementary Fig. 3a**). Using the CRISP-Cas9 approach, we generated stable knockout of IRF5 and PYK2 in these cells and validated their absence by western blot analysis (**Supplementary Fig. 3b**). After 5 days of ex vivo differentiation in the presence of GM-CSF, HoxB8 progenitors deficient in IRF5 or PYK2 gave rise to mature macrophages, comparable to the wt cells, but the levels of inflammatory cytokine and chemokine production were significantly reduced in HoxB8 macrophages deficient in IRF5 or PYK2 (**Supplementary Fig. 3c**). Thus, PYK2 acts as a critical regulator of IRF5-dependent transcription and inflammatory response induced by LPS in macrophages.

To characterise potential PYK2 target residues in IRF5, we employed phospho-proteomics. Endogenous IRF5 was immuno-precipitated from the lysates of LPS-stimulated WT and PYK2-deficient RAW264.7 macrophages, and the phospho-peptides were further enriched from the total proteolytic digests. Peptide masses and quantities were analysed by nano ultra-high-pressure liquid chromatography coupled with mass spectrometry (nUPLC-MS/MS). In line with previously published reports^14,15^ we identified the Ser-445 IKKβ-dependent site in both WT and PYK2-deficient cells (**Fig. 2f, Supplementary Fig. 4a, b**). In addition, endogenous IRF5 was phosphorylated at residues Ser-172, Ser-300, Tyr-334 in both WT and PYK2-deficient cells (**Fig. 2f, Supplementary Fig. 4a, b**). Interestingly, we could only detect Y171 phosphorylation in WT cells (**Fig. 2f, g Supplementary Fig. 4b**), while S56 and Y312 residues were modified in PYK2-deficient cells (**Fig. 2f, Supplementary Fig. 4a, b**), possibly reflecting on modification by other enzymes. In fact, Src tyrosine kinase Lyn was capable of phosphorylating IRF5 at orthologues of Y312 and Y334 sites in *in vitro* co-expression system^19^. We individually mutated these sites (Y171, Y312, Y334) as well as the published Y104 site^31^ into phenylalanine residues and explored the consequence of these mutations in the above-mentioned reporter and *in vitro* phosphorylation assays (**Supplementary Fig. 1a, f**). The Y172F mutant of human v2 IRF5 (Y171 of mouse IRF5) and the double mutant Y172, S173A (Y171, S172 of mouse IRF5) (**Fig. 2h**) both showed diminished ability to activate the TNF-luciferase reporter in the presence of PYK2, whereas the Y104F, Y329F (mouse Y312) and Y351F (mouse Y334) mutations had no inhibitory effect (**Fig. 2h**). Similarly, we observed reduction in PYK2-dependent phosphorylation of IRF5 Y172 and Y172/S173 but not other IRF5 mutants (**Fig. 2i**). Together, these results indicate that PYK2 mediates LPS-induced activation of IRF5, by phosphorylating the tyrosine site Y172 (mouse Y171) of IRF5.

Recently, specific inhibitors of PYK2 and a related kinase FAK have been developed^32^. One of them called defactinib (also known as VS-6063 or PF-04554878), with high selectivity to PYK2 and FAK1 and low affinity to kinases outside the family^33^, has been successfully used to tackle cancer in a mouse model and is currently being tested in clinical trials^34,35^. Here we explored the effect of defactinib on IRF5 activation and IRF5-dependent gene expression by macrophages. We first established the concentration range of defactinib well tolerated by RAW264.7 macrophages (**Supplementary Fig. 5a**). At such concentrations (0.3-1μM), it reduced LPS induced PYK2 phosphorylation (**Supplementary Fig. 5b**) and effectively inhibited TNF reporter activity and gene expression in RAW264.7 macrophages in a dose dependent manner (**Supplementary Fig. 5c, d)**. Moreover, defactinib inhibited TNF reporter activity in wt and IRF5-deficient RAW264.7 macrophages in which IRF5 expression was restored via ectopic expression of IRF5, but not in PYK2-deficient cells (**Fig. 3a**), suggesting that defactinib acts in an IRF5- and PYK2-specific manner. PF-573228 inhibitor with described 50 – 250-fold higher selectivity for FAK over PYK2^36^, also reduced TNF reporter activity but at a higher concentration than defactinib (**Supplementary Fig. 5e**). We speculated that PF-573228 may be targeting PYK2 in this system, and indeed observed no further inhibition in PYK2-deficient cells (**Supplementary Fig. 5e)**. The residual LPS-induced TNF reporter activity in cells treated with defactinib and PYK2 deficient cells (**Fig. 2c**) is consistent with the involvement of other PYK2 independent pathways in control of the TNF gene. As NF-κB is a known critical regulator of *TNF*^37^, we examined NF-kB activation in the cells deficient in PYK2 and/or treated with defactinib. Contrary to the results obtained in RAW264 cell lines stably expressing shRNA of PYK2^28^, we observed little effect on the p65/RelA phosphorylation and IkBa degradation in polyclonal RAW264 cell populations with CRISPR-Cas9 mediated knock-out of PYK2 and/or in cells treated with defactinib (**Supplementary Fig. 5f)**. We also detected no reduction in NF-kB reporter activity following treatment with defactinib (**Supplementary Fig. 5g)**. In addition, defactinib prevented LPS-induced nuclear translocation of IRF5 but not p65/RelA (**Fig. 3b**), ruling out a major role for NF-κB in PYK2 signalling pathway in these cells.

**Figure 3.**
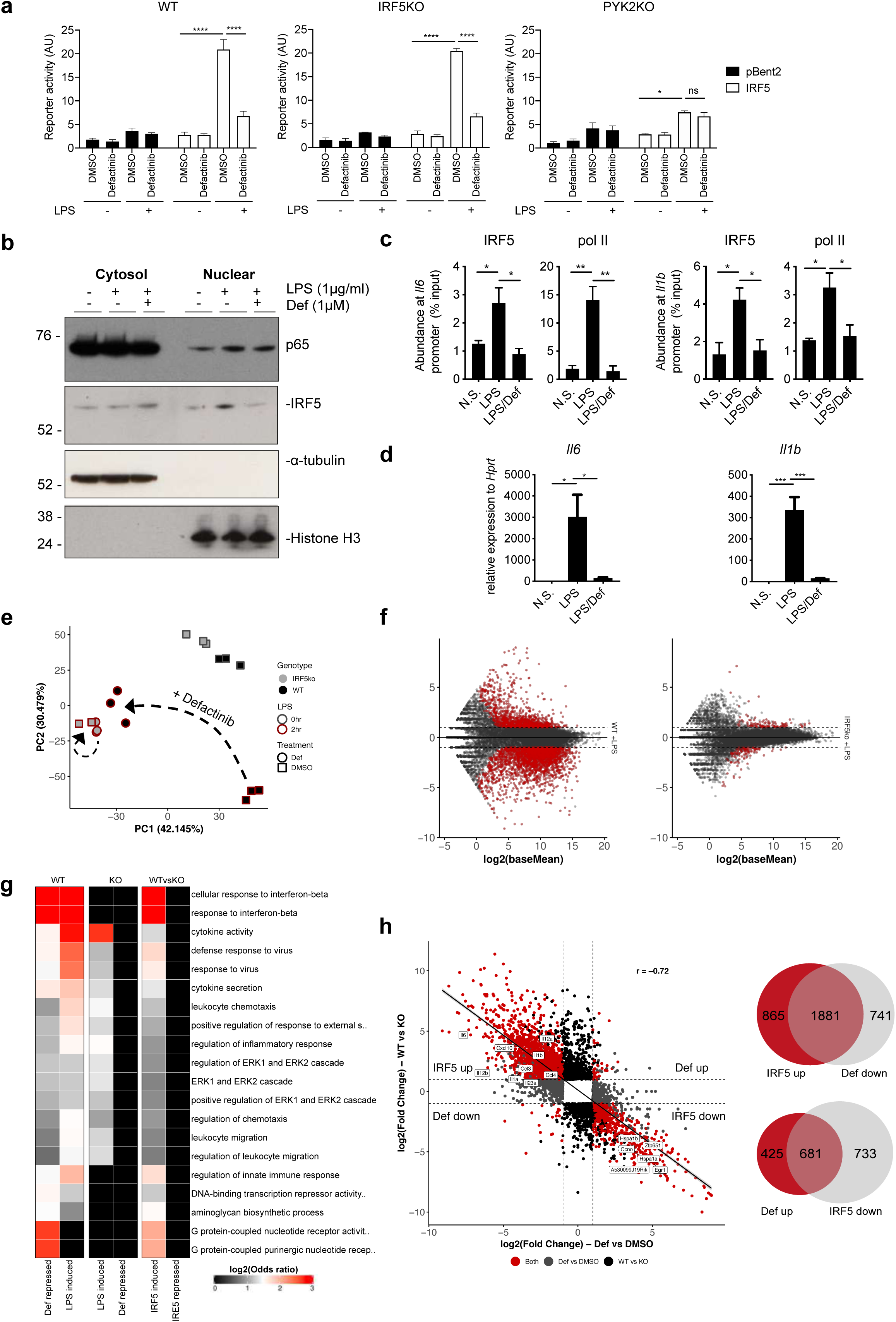
Inhibitors of PYK2 suppress IRF5-dependent gene induction. (**a**) TNF-luc reporter activity in the absence or presence of ectopically expressed IRF5 in wild type, IRF5 KO and PYK2 KO RAW264.7 cells pre-treated for 1 hr with 1 μM of defactinib (or DMSO control) and then stimulated with 1ug/ml of LPS for 6 hrs or left untreated. Data are shown as means +/− SEM from n=3 experiments. Comparison by two-way ANOVA \*\*\**P*<0.001 and \*\*\*\**P*<0.001. (**b**) RAW264.7 cells were fractionated into cytosolic and nuclear extracted following 1h pre-treatment with defactinib (1 μM) and 2 hr stimulation with LPS (1 μg/ml). (**c**) IRF5 and pol II binding to *Il6* and *Il1b* gene promoters in GM-CSF-differentiated mouse BMDMs pre-treated with 3.5 μM defactinib (def) or DMSO control and further stimulated with LPS for 2 hrs. Chromatin recruitment was analysed by ChIP. Data are normalized against chromatin amount in lysates (and expressed as percentage of input for each gene) and shown as mean values +/− SEM from n=3 individual mice, each performed in duplicates. Comparison by one-way ANOVA \**P*<0.05, \*\**P*<0.01 with multiple test corrections by Tukey. (**d**) *Il6* and *Il1b* expression levels in GM-BMDMs pre-treated with 3.5 μM defactinib (def) or DMSO control for 1 h, followed by LPS stimulation for 2 hrs. Data are shown as means +/−SEM for n=4 individual mice and analysed by one-way ANOVA \**P*<0.05 and \*\*\**P*<0.001. (**e**) PCA analysis of RNA-seq data from WT and Irf5^−/−^ pre-treated with 3.5 μM defactinib (def) or DMSO control for 1 h and further stimulated with LPS for 0 or 2 hrs. (**f**) MA plots depicting effect of defactinib on LPS stimulated BMDMs from WT or IRF5-/- mice. Differentially expressed genes (fold change > 1 and padj < 0.05) are highlighted in red. (**g**) GO enrichment analysis for differentially expressed genes (as in f). (**h**) Correlation analysis of IRF5 and defactinib regulated genes after 2 hrs LPS stimulation. Red indicates genes are differentially expressed (significance as in f) in both comparisons, genes regulated by IRF5 only (black), genes regulated by defactinib only (grey). Venn diagrams demonstrate overlap between IRF5 and defactinib regulated genes.

Mirroring our PYK2 deficiency data in RAW264.7 macrophages (**Fig. 2**), LPS-induced IRF5 and RNA polymerase II recruitment to its target promoters was suppressed in primary mouse bone marrow derived macrophages (BMDMs) treated with defactinib (**Fig. 3c**). The expression of some pro-inflammatory cytokines and chemokines was also effectively suppressed by defactinib (**Fig. 3d, Supplementary Fig. 6b**), without affecting cell viability (**Supplementary Fig. 6a**). Similar results were obtained in macrophages in response to activation of C-type lectin receptor Dectin-1 pathway, in which IRF5 has been shown to play a key role^38^, with WGP (dispersible whole glycan particles) (**Supplementary Fig. 6c**).

To investigate the global impact of PYK2 inhibition on IRF5 target gene expression, we compared LPS-induced transcriptomes in WT and IRF5 KO BMDMs treated with either defactinib or vehicle. Principle component analysis (PCA) of differentially expressed genes (DEGs) (p < 0.05) clearly separated WT and IRF5 KO, as well as untreated and LPS-treated sample groups (**Fig. 3e**). LPS stimulated WT samples treated with defactinib or vehicle were also clearly separated, with the defactinib treated samples grouping closely with the IRF5^−/−^. Conversely, defactinib had very little effect on IRF5^−/−^ cells stimulated with LPS (**Fig. 3e**). This was reflected in the number of DEGs: 4,026 for WT and only 217 for IRF5^−/−^ BMDMs treated with defactinib at 2 h of post LPS stimulation (**Fig. 3f, Supplementary Fig. 6d**). Gene ontology (GO) analysis for defactinib down-regulated genes revealed that they are predominantly pro-inflammatory in nature (e.g. cellular response to interferon-beta, regulation of inflammatory response, cytokine activity etc). These GO terms were also enriched in LPS induced genes, and in IRF5 up-regulated genes, suggesting that defactinib is highly specific for IRF5 target genes^22^ (**Fig. 3g**). We next investigated the correlation between IRF5 regulated genes and defactinib target genes. The majority of IRF5 up-regulated genes were strongly repressed by defactinib and there was a high degree of overlap between IRF5 up- and defactinib down-regulated genes, including *Il1a, Il1b, Il6, Il12a, Il12b, Il23a, Ccl3, Ccl4* etc (**Fig. 3h, Fig. Supplementary Fig. 6b**). Interestingly, there was also a smaller overlap between IRF5 down-regulated genes and defactinib up-regulated genes, suggesting that the actions of IRF5 and defactinib are in direct opposition to each other. Similar to our finding in mouse BMDMs, we saw robust inhibition of LPS-induced expression and production of IRF5-dependent cytokines in human monocyte derived macrophages treated with defactinib at 0.5-5 μM concentrations, which did not affect cell viability (**Supplementary Fig. 7a, b, c**). Taken together, PYK2 inhibition suppresses IRF5-dependent innate sensing and inflammatory response in both mouse and human macrophages.

IRF5 activity in mononuclear phagocytes (MNPs) plays a critical role in the pathogenesis of intestinal inflammation and that mice deficient in IRF5 are protected from overblown colitis^8,9^. Here we explored if inhibition of Pyk2 would also improve the intestinal immunopathology in a model of *Helicobacter hepaticus* infected and anti-IL-10R monoclonal antibodies administered (Hh+anti-IL10R) colitis, which is characterised by IL-23-dependent intestinal inflammation along with a robust T helper type 1/type 17 (Th1/Th17)-polarized effector T cell response^39^ (**Supplementary Fig. 8a**). As expected, Hh+anti-IL10R-infected mice developed inflammation in the colon after a week. However, intestinal pathology, as well as immune cell infiltrate, and PYK2 activation in colon tissue were reduced in the animals, which received defactinib (**Fig. 4a, b, c; Supplementary Fig. 8b, c**). We also observed attenuated induction of *Il1a, Il1b, Il6, Tnf, Il12b, Ccl4* and other pro-inflammatory cytokines and chemokines^40^ in the colon of defactinib-treated Hh+anti-IL-10R infected animals (**Fig. 4d**). To examine if the observed reduction in cytokine expression reflected on a lower number of monocytes infiltrating the colon or was related to their intrinsic reprogramming, we also analysed gene expression in total colonic leukocytes and isolated monocyte/macrophages. Interestingly, the downregulation of *Il6, Tnf, I12b* and *Ccl4* expression in response to defactinib was detected in (1) colon tissue, (2) total colonic leukocytes and (3) isolated colonic monocytes and macrophages, while the reduction of *Ccl5* expression and an upward trend in *Il10* expression was only observed in isolated macrophages, indicating that production of these mediators by other cells may mask the effect of IRF5 pathway inhibition in macrophages (**Supplementary Fig. 8d)**.

**Figure 4.**
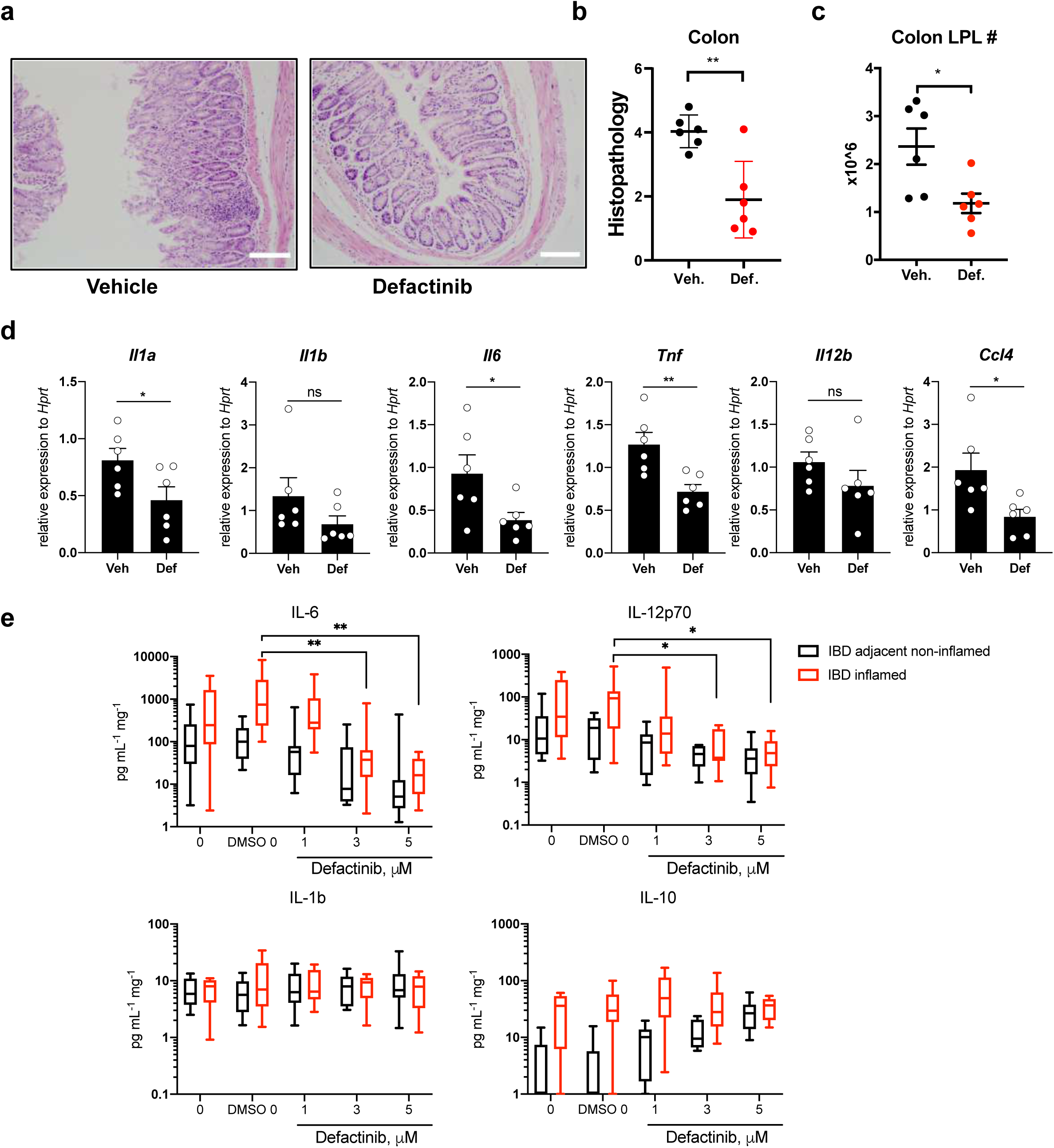
PYK2 inhibition reduced inflammation in Hh/anti-IL10R-model of murine colitis and in UC biopsies. (**a**) H&E staining of large intestine tissue sections, (**b**) histology scoring and (**c**) leukocyte content from *Hh*/anti-IL10R-treated mice, which received either vehicle or defactinib. **(d)** Cytokine/chemokine mRNA expression levels in mouse colon tissues from vehicle or defactinib Hh/anti-IL10R treated mice. Data from (b-d) are shown as means +/−SEM for n=6 mice \**P*<0.05 and \*\**P*<0.01 by unpaired Student’s t test. (**E**) Cytokine proteins levels in biopsies from ulcerative colitis patients from non-inflamed and inflamed tissues treated with defactinib at indicated concentrations per mg of tissue. Data are shown as means +/−SEM for n=10 human donors and analysed by two-way ANOVA where \**P*<0.05 and \*\**P*<0.01.

Next, we tested the impact of PYK2 inhibition in biopsies derived from the colonic mucosa of patients with active ulcerative colitis by measuring cytokine production at concentrations not affecting cell viability in these samples (**Supplementary Fig. 9 a, b**). We found elevated cytokine production in biopsies obtained from the sites of active inflammation in comparison to those from adjacent non-inflamed colon. Incubation with increasing doses of defactinib significantly lowered IL-6 and IL-12p70, identified as part of a cassette of inflammatory molecules that mark anti-TNFα-resistant IBD^41^ (**Fig. 4e**). We also observed a trend towards increased IL-10 production by the defactinib treated biopsies, but production of IL-1β appeared to be unaffected. Therefore, pharmacological inhibition of PYK2 effectively dampens intestinal inflammation, positioning defactinib and related PYK2 inhibitors as attractive molecules for repurposing to treat patients with UC.

In conclusion, we propose the following pathway involving Pyk2 and IRF5 in macrophages. Upon TLR4 stimulation by LPS or Dectin-1 stimulation by glucan, PYK2 is activated by phosphorylation at Tyr-402^28^. MyD88 is likely to be the essential linking adaptor between TLRs and PYK2/IRF5 complex as earlier studies have shown impairment of PYK2 activation in TLR ligand treated MyD88-deficient cells^28^. PYK2 autophosphorylation has been suggested to occur with the help of Src and possibly other kinases^42,43^. The recruitment of PYK2 to IRF5 upon Dectin-1 stimulation is likely to be Syk-dependent^44^. We show that PYK2 phosphorylates IRF5 on site Tyr-171 (mouse) contributing to its activation and transcription of pro-inflammatory cytokines (**Supplementary Fig. 10**). Multiple sites of phosphorylation detected in IRF5, both serine and tyrosine, highlight the complexity of IRF5 activation and multiplicity of signalling pathways^11,19,45,46^. Yet, a clear mechanistic interaction between two established IBD risk genes, PYK2 and IRF5, in macrophages, identified in this study, combined with an acceptable toxicological profile of PYK2 inhibitor defactinib shown in cancer clinical trials^47^, deserves a closer look from the therapeutic perspective. We propose that defactinib is an attractive molecules for repurposing to treat patients with ulcerative colitis^8,9^, and with other inflammatory conditions, such as arthritis^40,48^ acute lung injury^40^ and atherosclerosis^49^, in which IRF5 function in macrophages has been intimately linked to pathogenicity. It may even prove useful in dampening lung inflammation in the severe COVID 19 patients, whose lungs are filled with monocyte-derived macrophages expressing high levels of IRF molecules, including IRF5^50^.

## Acknowledgements

We are grateful to Dr Jelena Bezbradica-Mircovic (University of Oxford) for critical reading of the manuscript and useful comments. This work was supported by the Versus Arthritis (PhD studentship 209966 to HA), the BRC3 Gastroenterology and Mucosal Immunology (ALC and ST) and the Wellcome Trust (Investigator Award 209422/Z/17/Z to IAU). We thank GSK for providing their published kinase inhibitor library for this project.

## Methods

### Reagents

#### Animals

Mice were bred and maintained under SPF conditions in accredited animal facilities at the University of Oxford. All procedures were conducted according to the Operations of Animals in Scientific Procedures Act (ASPA) of 1986 and approved by the Kennedy Institute of Rheumatology Ethics Committee. Animals were housed in individually ventilated cages at a constant temperature with food and water ad libitum. C57Bl/6 mice were purchased from the University of Oxford BMS.

#### Cell culture

RAW264.7 and 293 TLR4/CD14/MD-2 cells were cultured in DMEM (Lonza) supplemented with 10 % FBS (Gibco) and 1 % Pen/Strep (Lonza). Bone marrow cells were extracted from wild type mice and cultured with recombinant GM-CSF (20ng/mL; Peprotech). On day 8, adherent cells were replated, and stimulated with either LPS (100ng/mL, Enzo) or whole glucan particles (100 μg/mL, Invivogen).

Human monocytes were isolated from leukocyte cones of healthy blood donors. Peripheral blood mononuclear cells (PBMC) were enriched by Ficoll gradient. Monocyte-derived macrophages were generated using adherence method selection and GM-CSF differentiation. Whole PBMC (50×10^6^) were plated in RPMI-1640 medium for 90 min. After 2 washes with PBS, adherent monocytes were differentiated into macrophages over a 5 days in the presence of 50 ng/mL GM-CSF (Peprotech) in RPMI supplemented with 10% foetal calf serum (FCS) (Sigma-Aldrich), 100 U/mL penicillin, 100 mg/mL streptomycin, 30 mM HEPES, and 0.05 mM β-mercaptoethanol.

Hoxb8 macrophage progenitors were a gift from the Sykes Lab (Harvard Medical School). Progenitors were cultured in RMPI-1640 medium (Lonza) supplemented with 10% FBS (Gibco), β-mercaptoethanol (30 mM; Life Technologies), recombinant GM-CSF (10ng/ml; Peprotec) and β-estradiol (1 μM; Sigma-Aldrich). To differentiate into macrophages, progenitors were washed three times with RPMI 1640 medium to remove the β-estradiol and incubated in complete RPMI 1640 medium supplemented with 10% heat-inactivated FBS, 30uM β-mercaptoethanol, and 20 ng/mL GM-CSF and incubated for 4 days. All cells were incubated in a 5 % CO_2_ humidified atmosphere at 37°C.

#### RNA extraction and Quantitative Real-Time PCR

Total RNAs were isolated from cells using RNeasy Mini Kit (Qiagen) and reverse transcribed to cDNA using High-Capacity cDNA Reverse Transcription Kit (Life Technologies) as per the manufacturer’s protocol. RNA from sorted cells was isolated utilising the RNeasy Micro kit (Qiagen). Real-time PCR reactions were performed on a ViiA7 system (Life Technologies) with Taqman primer sets for *Hprt, Ccl4, Ccl5, Il1a, Il1b, Il6, Il10, Il12a, Il12b, Il23a and Tnf*. Gene expression was analysed using the comparative Ct (ΔΔCt) method and normalised against *Hprt* levels or *RPLPO* levels for mouse or human, respectively.

#### RNA-Sequencing analysis

Libraries were sequenced on Illumina HiSeq4000 yielding > 40×10^6^ 150 b.p. paired end reads per sample. These were mapped to the mm10 genome using STAR^51^ with the options: “--runMode alignReads --outFilterMismatchNmax 2.” Uniquely mapped read pairs were counted over annotated genes using featureCounts^52^ with the options: “-T 18 -s 2 -Q 255.” Differential expression was then analysed with DESeq2^53^ and genes with fold changes > 2 and false discovery rates (FDRs) < 0.05 were deemed to be differentially expressed. Variance stabilised (VST) counts for all DESeq2 differentially expressed genes, likelihood ratio test, false discovery rates (FDRs) < 0.05, were used for dimensionality reduction. For direct comparisons genes with fold changes > 2 and FDR< 0.05 were deemed to be differentially expressed. Gene set enrichment analysis was performed using one-sided Fisher’s exact tests (as implemented in the ‘gsfisher’ R package https://github.com/sansomlab/gsfisher/). RNA sequencing data that support the findings of this study have been deposited in GEO with the accession code GSE141082.

#### Measurement of cytokine production

Mouse serum cytokine concentrations were analysed by ELISA (Mouse IL12p70, #DY419-05), and Cytometric Bead Array (IL6 #558301, IL1β #560232, BD Biosciences) as per manufacturer’s instructions. IL-1β, IL-6, and IL12p70 concentration in Human biopsy or cell culture supernatants was measured by Cytometric Bead Array (IL1β#558279, IL6# 558276, IL12p70#558283). IL-10 concentration in human intestinal biopsy supernatants was measured by ELISA (#DY217B-05, R&D systems). TNFα was measured in human monocyte-derived macrophage culture supernatants by ELISA (#DY210-05, R&D systems). All cytokine detection was performed according to manufacturer’s instructions.

#### Western blots

Cells were lysed in 1% TX-100 lysis buffer (1% v/v TX-100, 10% v/v glycerol, 1 mM EDTA, 150 mM NaCl, 50 mM Tris pH 7.8) supplemented with protease inhibitor cocktails (Roche). Lysates were incubated on ice for 30 min and cleared by centrifugation at 13,000 rpm for 10 min at 4°C. Protein quantification was performed with the Qubit assay (Thermo Fisher Scientific) according to the manufacturer’s protocol. 10 μg of lysates were boiled in Laemmli sample buffer (Bio-Rad), resolved on a NUPAGE 4-12% Bis-Tris gel (Invitrogen), and transferred onto a PVDF membrane (GE Healthcare) by wet western blotting. Membranes were blotted for antibodies for IRF5 (ab21689, Abcam), PYK2 (3292, CST), Phospho-Pyk2 Tyr402 (3291, CST), alpha-tubulin (3873, CST), Histone H3 (ab1791), GAPDH (ab9485, Abcam) and β-actin (A5441, Sigma), followed by HRP-conjugated secondary antibodies. Complexes were detected with the chemiluminescent substrate solution ECL (GE Healthcare).

#### Subcellular fractionation

Cell pellets were lysed in cytoplasmic lysis buffer (0.15 % NP-40, 10 mM Tris pH 7.5, 150 mM NaCl), incubated on ice for 10 minutes, and layered on top of cold sucrose buffer (10 mM Tris pH 7.5, 150 mM NaCl, 24 % w/v sucrose). The lysate was centrifuged at 13,000 rpm for 10 minutes at 4°C and the supernatant was collected as the cytosolic fraction. The nuclear pellet was lysed in RIPA buffer (150 mM NaCl, 1% NP-40. 0.5% Na-DOC, 0.1 % SDS, 50 mM Tris pH 8.0) and sonicated on the Biorupter sonicator (10 cycles of 30 seconds on/30 seconds off), followed by centrifugation at 13,000 rpm for 5 minutes at 4°C. The supernatant was collected as the nuclear fraction.

#### Immunoprecipitation

1×10^7^ million cells per immunoprecipitation were seeded and incubated overnight. Media was replaced with serum-free media for 1 hr, followed by LPS stimulation at indicated timepoints. Whole cell extracts were prepared with 1 % TX-100 lysis buffer as described above. Lysates were precleared with 100 μL TrueBlot Anti-Rabbit Ig IP beads (eBioscience) by rotating. Samples were incubated with 2 μg antibody for 2 hr, followed by 100 μL IP beads (50% slurry) by rotating overnight. Immunoprecipitates were washed three times with IP wash buffer (1 % NP-40, 150 mM NaCl, 1 mM EDTA, 20 mM Tris-HCl, pH 8) and eluted by boiling the samples for Laemmli sample buffer (Bio-Rad). Eluates were collected from the beads by centrifugation and resolved on a NUPAGE 4-12 % Bis-Tris gel (Invitrogen).

#### Generation of Pyk2 and IRF5 CRISPR knockouts

5,000 RAW264.7 cells/well were seeded in 96-well plates and infected the next day with PTK2B (ID:MM0000145196), and IRF5 (ID:MM0000200177) lentiviral particles (pLV-U6g-EPCG) provided by Sigma. Cells were transduced at a multiplicity of infection (MOI) of 10 in medium containing polybrene (8 μg mL^−1^) and spun at 1500 × g for 1 hr. After an overnight incubation, media was replaced with fresh media, and selected with 4 μg mL^−1^ puromycin (InvivoGen) for two weeks.

Hoxb8 macrophage progenitors were transduced with lentiCas9-v2 lentivirus targeting exon 2 of Irf5 (ID: ENSMUSG00000029771, gRNA ACCCTGGCGCCATGCCACGAGG) and exon3 of PYK2 (ID: ENSMUSG00000059456, gRNA CCCTATTCGCCCACTCAGG). The lentiCas9-v2 plasmid was a gift from Feng Zhang (Addgene plasmid #52961). Briefly, the lentiCas9-v2 lentivirus were produced from HEK-293FT cells transfected with the lentiCas9-v2 plasmid mixed at a 2:1:1 DNA ratio of the lentiviral packaging plasmids pMD2.G (Addgene plasmid #12259) and psPAX2 (Addgene plasmid #12260) at a 2:1:1 ratio. Media was replaced 16 hours post-transfection. Two days post transfection, the lentivirus containing mediums were harvested, filtered and added onto Hoxb8 macrophage progenitor cells at a final concentration of 8ug/ml polybrene. Transduced cells were allowed to grow for additional four days and selected with 6ug/ml Puromycin for the targeted knockout of Irf5 and Pyk2.

#### Chromatin Immunoprecipitation

1×10^7^ million cells per ChIP were seeded and incubated overnight. GM-BMDMs cells were pretreated with defactinib or DMSO vehicle for 1 hour, followed by LPS (100 ng/ml) for 2 hours. RAW264.7 cells were stimulated with LPS (500 ng/ml) for 2 hours. Cells were fixed in formaldehyde, quenched with Tris pH 7.5 and washed in PBS. Nuclear lysates were isolated as previously described^22^ and sonicated with a Bioruptor (Diagenode) for 8 cycles (GM-BMDMs) or 10 cycles (RAW264.7). Lysates were immunoprecipitated with 5 μg of anti-IRF5 (ab21689; Abcam), anti-RNA Polymerase II (MMS-128P; Biolegend), or Rabbit Anti-Mouse IgG (ab46540; Abcam). Immunoprecipitated DNA was purified with the PCR Purification Kit (Qiagen). qPCR analysis was carried out in duplicates and represented as % input. Primer sequences *Il1a* (ACTTCTGGTGCTCATCTGTCATGTT, GCTCTATGGTTCCTGTGTCTGTAGG), *Il1b* (GGATGTGCGGAACAAAGGTAGGCACG, ACTCCAACTGCAAAGCTCCCTCAGC), *Il6* (GAGAGAGGAGTGTGAGGCAGAGAGC, GGTTGTCACCAGCATCAGTCCCAAG), *Il12b* (GCAAGGTAAGTTCTCTCCTCTTCCC, AATGACTATTTGAAGCCCCTGTCGT), TNF (GCTAAGTTCTTCCCCATGGATGTCCC, ACCCATTTCTTCTCTGTCCTCCAGAGC).

#### Kinase inhibitors screening and luciferase reporter assay

RAW264.7 cells were seeded in eighteen 96-well plates at 50,000 cells/well a day before transfection as described above. 1hr prior to LPS treatment, cells were treated with 10uM of inhibitors in quadruplicates. For experiment wells (n=4 for each inhibitor set AG-AK, total amounts for 16 plates (AG-AJ1-4) – 80 wells per plate and 2 plates (AK1,2-3,4)– 96 wells per plate) 21 ml of Opti-Mem (Gibco) was mixed with DNA: 50 μg of pBent-HA-IRF5, 50 μg of pGL3-5’3TNF-luc and 25 μg of pEAK8-Renilla, vectors described in ^21,54^. 5ml of Opti-Mem mixed with 200 μl of Plus reagent was added to the DNA solution and incubated for 5-15 min. Then, 5ml of Opti-Mem was mixed with 500 μl of Lipofectamine LTX reagent, added to the DNA/Plus solution and incubated for 30 min. For controls (amount for 4 plates – two 4 well-rows each), 800 μl of Opti-Mem was mixed with DNA: 2 μg of pBent-HA-IRF5 or pBent2-empty, 2 μg of pGL3-5’3’-TNF-luc and 1 μg of pEAK8-Renilla. 200 μl of Opti-Mem mixed with 8 μl of Plus reagent was added to the DNA solution and incubated for 5-15 min. Then, 200 μl of Opti-Mem was mixed with 20 μl of Lipofectamine LTX reagent, added to the DNA/Plus solution and incubated for 30 min. To transfect cells, 20 μl of the DNA/transfection reagent mix was added per well. Next day, cells were pre-incubated for 1 hr with 20 μl of inhibitor (or 1% DMSO to control wells) in serum-free DMEM (final conc. 0.1, 1 or 10 μM in 1% DMSO). Then, 1μg/ml of LPS was added to the cells and 6 hours later the culture medium was and the plate-bound cells were kept frozen (−20 C). Cells were lysed using the Dual-Glo Luciferase Assay kit (Promega) according to the manufacturer’s protocol and analysed in a FLUOstar Omega microplate reader (BMG Labtech). Raw firefly luciferase activities (or values normalized against Renilla luciferase activities) in wells incubated with the kinase inhibitors were divided by the luciferase activity values in the control wells (DMSO vehicle only, cells expressing IRF5 and stimulated with LPS) and expressed as part of a whole or a percentage of IRF5 reporter activity (which was 1 or 100% in cells treated with DMSO only).

### Kinase assays

293 ET cells were plated at 250,000 cells/well in six-well plates and a day later were transfected with 1 μg of pBent2-HA-IRF5 and 1 μg of plasmid encoding one of the myc-tagged candidate IRF5 kinases (in the pEAK8-myc vector) using Lipofectamine2000™ (Life technologies) according to the manufacturers protocol. The cell lysates were subjected to kinase assays using a modification of an established protocol^55^. Cells were washed in PBS and lysed on the ice in kinase reaction buffer (20 mM HEPES pH 7.5, 137 mM NaCl, 0.5 mM EGTA, 25 mM MgCl2, 0.2% Triton X-100, 10% Glycerol) with added protease (EDTA-free! complete-mini protease inhibitor cocktail, Roche, #11836170001) and phosphatase inhibitors (phosphatase inhibitor cocktail II, Sigma, #P5726). 2 mM of TCEP (#C4706, Sigma), 1mM of GTP (#G8877, Sigma) and 50 μM S-γ-ATP (ab138911, Abcam) was added sequentially to lysing samples. The reactions were mixed by vortexing and kept an a rocking surface for 1h at 37C. The reactions were stopped by adding 50 mM EDTA and moving them on ice. p-Nitrobenzyl mesylate (PNBM, ab138910, Abcam) was dissolved in DMSO to 50 mM. PNBM working solution was prepared by adding 5 μL of deionized water five times mixing after each addition to 25 μL of the PNBM stock, and then was added 1/10 to kinase reactions (at 2.5 mM), which were further incubated for 2 hrs at room temperature. The bulk of the reactions were subjected to immunoprecipitations using anti-thiophosphate-ester antibody (1 μg per reaction, ab92570, Abcam) and Protein G Sepharose 4 Fast Flow Media (#11524935, GE Healthcare) to pull-down phosphorylated IRF5. The rest of the reactions and the pull-downs were mixed with SDS PAGE loading buffer and subjected to SDS PAGE following Western blotting.

#### Mass spectrometry analysis

IRF5 was immunoprecipitated from LPS-stimulated WT and PYK2 KO RAW264.7 cells (5×10^7^ cells) as described in immunoprecipitation section. Eluents were subjected to in-solution digestion as described previously^56^. In brief samples were reduced and alkylated before double precipitation with Chloroform/Methanol as described^57^. Protein pellets were resuspended in 50 µL 6M urea for solubilisation. The samples were diluted to 1M Urea in 100mM Tris buffer for tryptic digest. Following overnight digestion, peptides were acidified with 3% Formic acid and desalted with solid phase extraction Sola cartridges (Thermo). Peptides were eluted with 600 uL glycolic acid solution (1M glycolic acid, 80% acetonitrile, 5% trifluoroacetic acid). Phospho-peptide enrichment was performed using a TiO_2_ protocol as described ^58^ with eluates from the Sola cartridges adjusted to 1 mL with 1M glycolic acid solution and incubated for 5 minutes with 50 uL TiO_2_ bead slurry solution. Bead washes (200 uL) were carried out as previously described. In short, beads were sequentially washed with 200 uL glycolic acid solution, ammonium acetate solution (100 mM ammonium acetate in 25% acetonitrile) and 10% acetonitrile solution repeated in triplicate. Phospho-peptides were eluted, following incubation for 5 minutes at room temperature with 50 ul ammonia solution (5%) and centrifuged, this was repeated in triplicate. The three eluate fractions were combined and dried using a SpeedVac and pellets were stored at −80°C until analysis. For analysis by nano-liquid chromatography tandem mass spectrometry (nLC-MS/MS), a Dionex UHPLC system coupled to an Orbitrap Fusion Lumos mass spectrometer was used as described previously ^59^. Raw MS files were subjected to processing using PEAKS (version 8.5) software and searched against the UniProtSP Mus Musculus database. Searches included the data refine, denovo PEAKS and PEAKS PTM modes, the latter of which included phosphorylation on Ser (S), Thr (T) and Tyr (Y) residues. The proteomics data and MS raw files have been deposited to Proteome Xchange Consortium via the PRIDE^60^ partner repository with the dataset identifier PXD014033 (https://www.ebi.ac.uk/pride/archive/).

#### *Helicobacter hepaticus*-induced colitis model

Mice were free of known intestinal pathogens and negative for Helicobacter species. Animals from each experimental group were cohoused. On days 0, 1, and 2, mice were injected i.v. with 1 mg/kg defactinib or vehicle (5% DMSO, 2.5% Solutol HS (Sigma), 2.5% absolute ethanol, 90% Dulbecco’s PBS). Daily, starting from day 3, mice were injected i.p. with 5 mg/kg defactinib or vehicle. 30 minutes after the initial i.v. injection, mice were infected with 1×10^8^ colony forming units Hh on days 0 and 1 by oral gavage with a 22G curved, blunted needle (Popper & Sons). Mice were injected intraperitoneally once on day 0 with 1 mg anti-IL10R blocking antibody (clone 1B1.2). Infected mice were monitored daily for colitis symptoms. Mice were culled one week after day of infection, and organs were harvested for analysis.

#### Isolation of lamina propria leukocytes

Colons and/or caeca were harvested from mice, washed in PBS/BSA and content flushed with forceps. Intestines were then opened longitudinally and washed once more before blotting to remove mucus. Gut tissue was then cut into 1 cm long pieces and placed in 50 mL centrifuge tube (Greiner) in ice cold PBS + 0.1% BSA. Colons were incubated 2 times at 200 rpm in 40 mL HBSS + 0.1% BSA + 1% Penicillin-Streptomycin (PS, Lonza) + 5mM EDTA (Sigma-Aldrich) at 37 °C for 10 min before the supernatant was aspirated. Tissue was placed in 40 mL PBS + 0.1% BSA + 1% PS for 5 min. Intestines were then incubated with 20 mL RPMI + 10% FCS +1% PS + 2.5 U/mL Collagenase VIII (Sigma-Aldrich) + 2 U/mL DNAse I (Roche), shaking at 200 rpm for 45 mins – 1 hour at 37 °C. Supernatant was filtered through a 70 μm cell strainer to which 30 mL of ice cold PBS + 0.1% BSA + 1% PS + 5 mM EDTA was added to ablate collagenase/DNase activity. Cells were washed in 30 mL PBS/BSA before filtering once more through a 40 μm cell strainer. The cells were then pelleted by centrifugation at 400 rcf for 10 minutes at 4 °C and resuspended in 1 mL RPMI + 10% FCS + 1% PS before counting.

#### Flow cytometry

CBA quantification of cytokine levels were performed on a FACSCanto II (BD) and analysed using Flowjo (Treestar Inc.). Acquisition of mouse samples was performed using either LSR II or Fortessa X20 flow cytometers with FACSDiva (BD), followed by analysis in Flowjo (Treestar Inc.). Gating strategy in **Fig. S8e**.

#### Extracellular labelling of cells

5×10^5^ – 2×10^6^ cells were plated on U-bottom 96 well plates. Cells were washed twice with 150 μL FACS buffer (PBS + 0.1 % BSA + 1 mM EDTA + 0.01% Sodium Azide) at 400 rcf for 3 min 4°C. Cells were then Fc blocked for 10 min with αCD16/CD32 (BD) 1/100 in 20 μL FACS buffer at room temperature (RT) followed by washing once in 150 μL FACS buffer. Fixable Viability Dye eFluor®780 (ThermoFisher) and primary extracellular antibodies (Table 1.1) were added for 30 min at 4 °C in 20 μL FACS buffer in the dark. Labelled cells were then washed twice with 150 μL FACS buffer. Cells were then fixed for 30 mins in 50 μL Cytofix (BD), washed twice with 150 μL FACS buffer, and resuspended in 200 μL FACS buffer before acquisition.

**Table 1.1.** Antibodies used in flow cytometry analysis.

| **Table.1.1 Antibodies used in flow cytometry analysis** |  |
| --- | --- |
| **Antigen** | **Clone** |
| CD45 | 30-F11 |
| CD11b | M1/70 |
| CD11c | N418 |
| Siglec F | E50-2440 |
| Ly6G | 1A8 |
| F4/80 | BM8 |
| CD103 | 2E7 |
| CD206 | CO68L2 |
| MHC II | M5/114.15.2 |
| CD19 | 6D5 |
| CD138 | 281-2 |
| NK1.1 | PK136 |
| CD3e | 145-2C11 |
| TCRgd | GL3 |
| Ter119 | TER-119 |

#### FACS sorting

Colon lamina propria cells were prepared as described above. A small aliquot of each sample was stored in RNAlater (Sigma Aldrich) for further processing. Two to three samples were pooled in order to gain sufficient numbers for sorting. The cells were labelled as described above with the antibodies in Table 1.1, except no fixation step was performed. Labelled cells were washed twice with 1mL FACS buffer and resuspended in 500uL FACS buffer containing DNAse I (10ug/mL, Roche). Sorting of the cells was performed into 500uL RNAlater on FACSAria III (BD Biosciences) at the Kennedy Institute of Rheumatology FACS facility.

#### Culture of UC patient colonic mucosal biopsies

Intestinal pinch biopsies were obtained from Ulcerative Colitis patients registered in the Oxford IBD Cohort, attending the John Radcliffe Hospital Gastroenterology Unit (Oxford, UK). This cohort comprises 1896 patients with UC, median age 31 at diagnosis, treated with biological therapy (23%) or conventional steroids/immunomodulators (77%) for active disease, in addition to mesalazine. Biopsies were collected during routine endoscopy. Informed, written consent was obtained from all donors. Human experimental protocols were approved by the NHS Research Ethics System (Reference numbers: 16/YH/0247). Biopsies were washed in PBS and transferred into wells containing RPMI-1640 + 10% FCS + 20 μg/mL G418 (Thermo Fisher) + 20 U/mL Pen/Strep and cultured for 24 hours.

#### UC biopsy viability assessment

Biopsies were fixed in 4% PFA in PBS (#30525-89-4, Santa Cruz) for 24 hrs at RT and transferred to 70% ethanol. Fixed biopsies were then dehydrated and embedded in paraffin blocks, and 5 μm sections were cut. Embedding and sectioning of tissues was carried out by the Kennedy Institute of Rheumatology Histology Facility (University of Oxford). Viability of intestinal biopsies was measured by TACS® TdT *in situ* (Fluorescein) TUNEL assay (#4812-30-K, R&D systems) according to manufacturer’s instructions. Sections were then mounted in Glycerol Mounting Medium with DAPI and DABCO (#ab188804, Abcam) and cover-slipped. Images of three non-sequential sections per sample were acquired. Three images per section were acquired at 20x magnification using a BX51 microscope (Olympus). To generate the apoptotic index, the total cell number was enumerated by counting DAPI^+^, and TUNEL(FITC)^+^ cells in ImageJ, and calculating the percentage of total cells that were TUNEL^+^.

#### Histopathological assessment

Post-sacrifice, 0.5 cm pieces of caecum, and proximal, mid and distal colon were fixed in PBS + 4% paraformaldehyde (Sigma Aldrich). Fixed tissue was embedded in paraffin blocks, and sectioned using a microtome and stained with Haematoxylin and Eosin (H&E) by the Kennedy Institute of Rheumatology Histology Facility (Kennedy Institute of Rheumatology, University of Oxford). Sections were scored in a blinded manner by two researchers according to ^61^.

#### Cell viability

50,000 cells/well were seeded and incubated overnight. Cell viability was assessed using the Promega CellTiter-Glo® Luminescent kit per the manufacturers protocol and luminescence was measured in a FLUOstar Omega microplate reader (BMG Labtech). Samples were tested in triplicate and normalised to untreated wells.

#### Protein isolation from colon tissue

1.5 ml Bioruptor Microtubes were filled with 250 mg of Protein Extraction Beads (Diagenode) and filled with RIPA buffer (supplemented with protease and phosphatase inhibitors). 10 mg of tissue was added to the tubes and vortexed briefly. Tubes were sonicated on the Biorupter Pico with 30 sec ON/30 sec OFF for 6 cycles at 4°C. After each 2 cycles, tubes were vortexed. The supernatant was transferred to a new tube and centrifuged at 13,000 rpm for 10 minutes at 4°C. The supernatant was transferred to a new tube and 80 μg of lysate was used for immunoblot analysis.

## Supporting information

**Source Data 1**. Reporter gene assays screening of GSK PKIS set for IRF5 activation inhibitors.

**Supplementary Figure 1. Screening and validation assays to identify novel IRF5 kinases** TNF and ISRE-luc reporter activity in (**a**) HEK-TLR4 and (**b**) RAW264.7 cells co-expressing IRF5 or empty pBent2 plasmid control. Cells were stimulated with LPS (1μg/ml) for 6 hrs or left untreated. Data are shown as means +/− SEM for 3 independent experiments each performed in triplicates and analysed by two-way ANOVA where \**P*<0.05 and \*\*\*\**P*<0.0001. (**c**) A scheme of small molecule screening for candidate IRF5 kinases. RAW cells were transfected with plasmids encoding for IRF5 and TNF-luciferase reporter as well as constitutively expressed Renilla luciferase. (1) 24 hrs after transfection cells were pre-treated with a library of inhibitors (four replicate wells per inhibitor) for 1hr (2) and stimulated with 1 μg/mL of LPS for 6 hrs (3) before lysing cells for luminometry. (**d**) Stratification of the kinase inhibitors used in the screen based on the degree of IRF5 reporter inhibition. The numbers are shown based on activities of the firefly luciferase reporter, raw values or normalised to Renilla luciferase activities to account for non-specific impact of cell viability. Out of 365 molecules, 57 inhibited IRF5 reporter activity by 30-50%, 34 – 50-70%, 8 – 70-80%, 5 – 80-90% and 4 by >90% (the normalised activity readout). (**e**) Activities of top 10 IRF5 reporter inhibitors are shown where the dataset was analysed based on raw firefly luciferase or normalised to Renilla values. The compounds indicated with numbers are in top 10 independently of normalisation. To calculate reporter activity luciferase values (raw or normalised to Renilla) in wells incubated with kinase inhibitors were divided by the luciferase activity values in the control wells (DMSO vehicle only, cells expressing IRF5 and stimulated with LPS). (**f**) A scheme of a modified-ATP based IRF5 kinase assay. Cells co-expressing HA-tagged IRF5 with either of the candidate kinases were lysed and incubated with S-γ-ATP. The newly-produced phosphate groups were further labelled using a reaction with PNBM and the modified proteins were pulled down using anti-thiophosphate ester antibody. (**g**) Table summarising functional validation of candidate kinases related to Fig. 1b-d.

**Supplementary Figure 2. PYK2 regulates IRF5 activation and IRF5-mediated transcription**.

(**a**) Western blot analysis of PYK2 and IRF5 expression in RAW 264.7 cells transfected with control, PYK2 or IRF5 CRISPR-based knockout constructs. (**b)** Immunoblot analysis for restoring PYK2 expression in PYK2 deficient RAW264.7 cells. **(c)** TNF-luciferase reported activity in WT and PYK2 KO RAW264.7 cells co-transfected with pBent2-Empty, HA-IRF5, or Myc-PYK2 along with TNF-firefly Luc and pRLTK-Renilla Luc. Cells were stimulated with LPS (1 μg/ml) or left untreated for a further 6 hrs. (**d**) IRF5 and pol II binding to *Tnf* gene promoter in resting or LPS-treated (2h, 500 ng/ml) wild type or PYK2 KO RAW264.7 cells as measured by the chromatin immunoprecipitation (ChIP) method. A non-specific IgG antibody was used as a negative control for ChIP. Data are normalized against chromatin amount in lysates (and expressed as percentage of input for each gene). (**e**) Gene expression levels in wild type, PYK2 KO or IRF5 KO RAW264.7 cells stimulated with LPS (500 ng/ml) for 0, 2, or 4 hrs. Gene expression was measured by qPCR. All values in (c-e) are shown as mean values +/− SEM from n=3 experiments. Comparison by two-way ANOVA \**P*<0.05, ** *P*<0.01, *** *P*<0.001, and **** *P*<0.0001.

**Supplementary Figure 3. PYK2 regulates IRF5-mediated transcription in macrophages**.

(**a**) Flow cytometry analysis of HoxB8 cells after 5 days of differentiation with GM-CSF. Day 0 corresponds to cells prior to differentiation. **(b)** Western blot analysis of IRF5 and PYK2 expression in Hoxb8 macrophage progenitors transfected with PYK2 or IRF5 CRISPR-based knockout constructs. **(c)** Gene expression levels in wild type, PYK2 KO or IRF5 KO HoxB8 macrophages stimulated with LPS (100 ng/ml) 2 hrs. Gene expression was measured by qPCR. Values shown as mean values +/− SEM from n=3 experiments. Comparison by two-way ANOVA \**P*<0.05, ** *P*<0.01, *** *P*<0.001, and **** *P*<0.0001.

**Supplementary Figure 4. Mass spectrometry analysis detects PYK2-dependent phosphorylated sites in IRF5**.

(**a**) MS/MS spectra indicating mouse IRF5 S300, Y334, S445, S56, Y312, S172 phosphorylation in LPS-stimulated WT and PYK2 KO cells. Fragmentation ions of the b- and y-series are indicated in blue and red, respectively. **(b)** Identification of endogenous IRF5 phosphorylation sites by tandem mass spectrometry (MS/MS) in LPS-stimulated WT (top panel) and PYK2 KO (lower panel) RAW264.7 cells in which the peptides identified by LC-MS/MS are underlined by a blue line. Each line corresponds to a unique MS/MS spectrum. Post-translational modifications including cysteine carbamidomethylation (orange box) and phosphorylation (red box) are indicated. The location of phosphorylated Tyr 171 (Y171) and Ser172 (S172) observed in WT IRF5 are marked in a red box. (**c**) IRF5 sites in mouse and equivalent position in human isoform 2.

**Supplementary Figure 5. Defactinib affects Pyk2 phosphorylation and IRF5-dependent gene expression at concentrations that do not affect cell viability**.

(**a**) Cell viability in RAW264.7 cells pre-treated with DMSO/Defactinib for 1 hr followed by LPS (1 μg/mL) for 6 hrs. IC_50_, inhibitor concentration at which 50% decline in cell viability was observed compared to control (DMSO). (**b**) Immunoblot of lysates of RAW264.7 cells pretreated for 1 h with 1 μM defactinib (def) or DMSO control, and stimulated with LPS (1 μg/mL) for 30 min. Blots were probed with Abs specific for PYK2 phosphorylated on Tyr-402, total PYK2 and GAPDH. (**c**) TNF-luc reporter activity in the absence or presence of ectopically expressed IRF5 in RAW264.7 cells pre-treated for 1 hr with defactinib (or DMSO control) at indicated concentrations followed by LPS (1ug/ml) for 6 hrs. (**d**) Gene expression levels in RAW264.7 cell pre-treated with defactinib (def) or DMSO control for 1 h, followed by LPS stimulation for 4 hrs. Data are shown as means +/−SEM for n=3 and analysed by one-way ANOVA. (**e**) TNF-luc reporter activity in the absence or presence of ectopically expressed IRF5 in wild type, IRF5 KO and PYK2 KO RAW264.7 cells pre-treated for 1 hr with defactinib (Def), PF-573228 (PF) inhibitor or DMSO control at indicated concentrations and stimulated with LPS (1 μg/ml) for a further 6 hrs. (**f**) WT and PYK2 KO RAW264.7 cells were pre-treated with 1 μM of defactinib (Def) or DMSO vehicle control for 1 h followed by LPS (500 ng/ml) at indicated timepoints. Cell lysates were subjected for immunoblot using indicated antibodies. (**g**) NFkB-luc reporter activity in HEK-TLR4 cells co-expressing IRF5 or empty pBent2 plasmid control. Cells were pretreated with defactinib (1μM) or DMSO control for 1 hr and stimulated with LPS (1μg/ml) for 6 hrs. Data for (c), (e) and (g) are shown as means +/− SEM of three independent experiments, and analysed by 2 way ANOVA. \*\*\**P*<0.001 and \*\*\*\**P*<0.0001.

**Supplementary Figure 6. Defactinib affects IRF5-dependent gene expression**

**(a)** Cell viability in GM-BMDMs pre-treated with DMSO/Defactinib for 1 hr followed by LPS (100 ng/ml) for 2 hrs. **(b)** Gene expression levels in GM-BMDMs pre-treated with 3.5 μM defactinib (def) or DMSO control for 1 h, followed by LPS (100 ng/ml) or **(c)** WGP (100 ug/ml) for 2 hrs. Data are shown as means +/−SEM for n=4 individual mice and analysed by one-way ANOVA \**P*<0.05, \*\**P*<0.01, and ***P<0.001. **(d)** Number of DE genes from RNA-seq.

**Supplementary Figure 7. Defactinib affects gene expression in human monocyte-derived macrophages**.

**(a**) Cell viability was measured in human monocyte-derived macrophages (hMDMs) after 3hrs or 24 hrs of treatment with defactinib. (**b**) Cytokine mRNA expression levels in human monocyte-derived macrophages pre-treated with defactinib (def, 5 μM) for 1 h followed by LPS stimulation (100 ng/ml) for 2 h. Data are shown as means +/−SEM for n=4 and analysed by one-way ANOVA where **P<0.01 and ***P<0.001. (**c**) Cytokine proteins levels in human monocyte-derived macrophages pre-treated with various amounts of defactinib for 1h, followed by stimulation with LPS for 24 h.

**Supplementary Figure 8. PYK2 inhibition in Hh/anti-IL10R-model of murine colitis**.

**(a**) Defactinib treatment regime during the initiation phase of mouse Hh+anti-IL-10R colitis. **(b**) Immune cell infiltrate from Hh/antil-IL10R-treated mice, which received either vehicle or defactinib. (**c**) PYK2 autophosphorylation (pY402) in vehicle or defactinib treated mice (6 mice each) assessed by western blot analysis. Lysates from LPS stimulated GM-BMDMs included as a positive control. (**d**) Gene expression levels in colon tissue, leukocytes, monocytes/macrophages from vehicle or defactinib Hh/anti-IL10R treated mice. (**e**) Gating strategy for (b). Data in (b) and (d) are shown as means +/−SEM. \**P*<0.05 and \*\**P*<0.01 by unpaired Student t test.

**Supplementary Figure 9. PYK2 inhibition in UC biopsies**.

**(a)** Defactinib treatment of human biopsies from inflamed and non-inflamed sites of patients with ulcerative colitis. (**b**) Cell viability by TUNEL assay was measured in colon biopsies after 24 hr treatment with defactinib.

**Supplementary Figure 10. Proposed model of IRF5 activation by PYK2 in macrophages**. LPS stimulation of TLR4 leads to PYK2 autophosphorylation on site Y402. Activated PYK2 phosphorylates IRF5 at site Y171 (mouse). IRF5 translocates to the nucleus and activates target genes. Serine kinases IRAK4, TAK1, and IKKβ have been proposed to phosphorylate and activate IRF5 downstream of the TLR-MyD88 pathway^12–15^, while IKKα and Lyn negatively regulated IRF5^19,62^. Dectin-1 stimulation by whole glucan particles also leads to IRF5 mediated transcription and is likely to be Syk-dependent.

