## Supplementary Fig. 1 for "PYK2 controls intestinal inflammation via activation of IRF5 in macrophages"

### Supplementary Figure S1

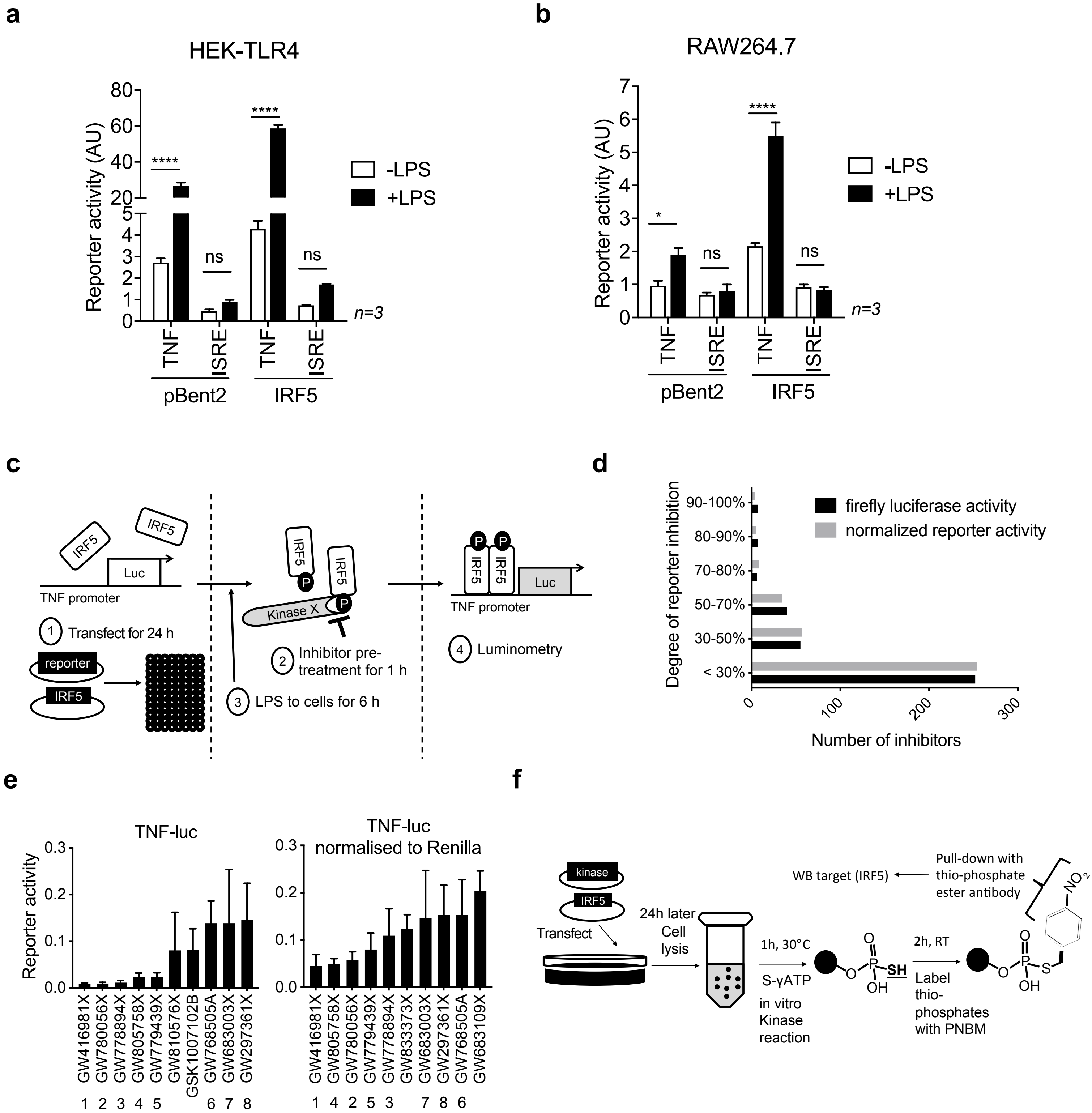


g

| | KINASE | | | | | | |
|---|---|---|---|---|---|---|---|
| | ARK5 | CLK2 | HIPK4 | JNK2 | MARK3 | MST1 | PYK2 |
| TNF-luciferase (overexpression) | X | ✓ | ✓ | ✓ | X | ✓ | ✓ |
| IRF5 binding (overexpression) | ✓ | X | ✓ | ✓ | X | X | ✓ |
| *In vitro* kinase assay (overexpression) | X | X | X | ✓ | ✓ | ✓ | ✓ |
| IBD locus | X | X | X | X | X | X | ✓ |
