## Supplementary figures and images for "PYK2 controls intestinal inflammation via activation of IRF5 in macrophages"

### Supplementary Fig. 2

#### Supplementary Figure S2

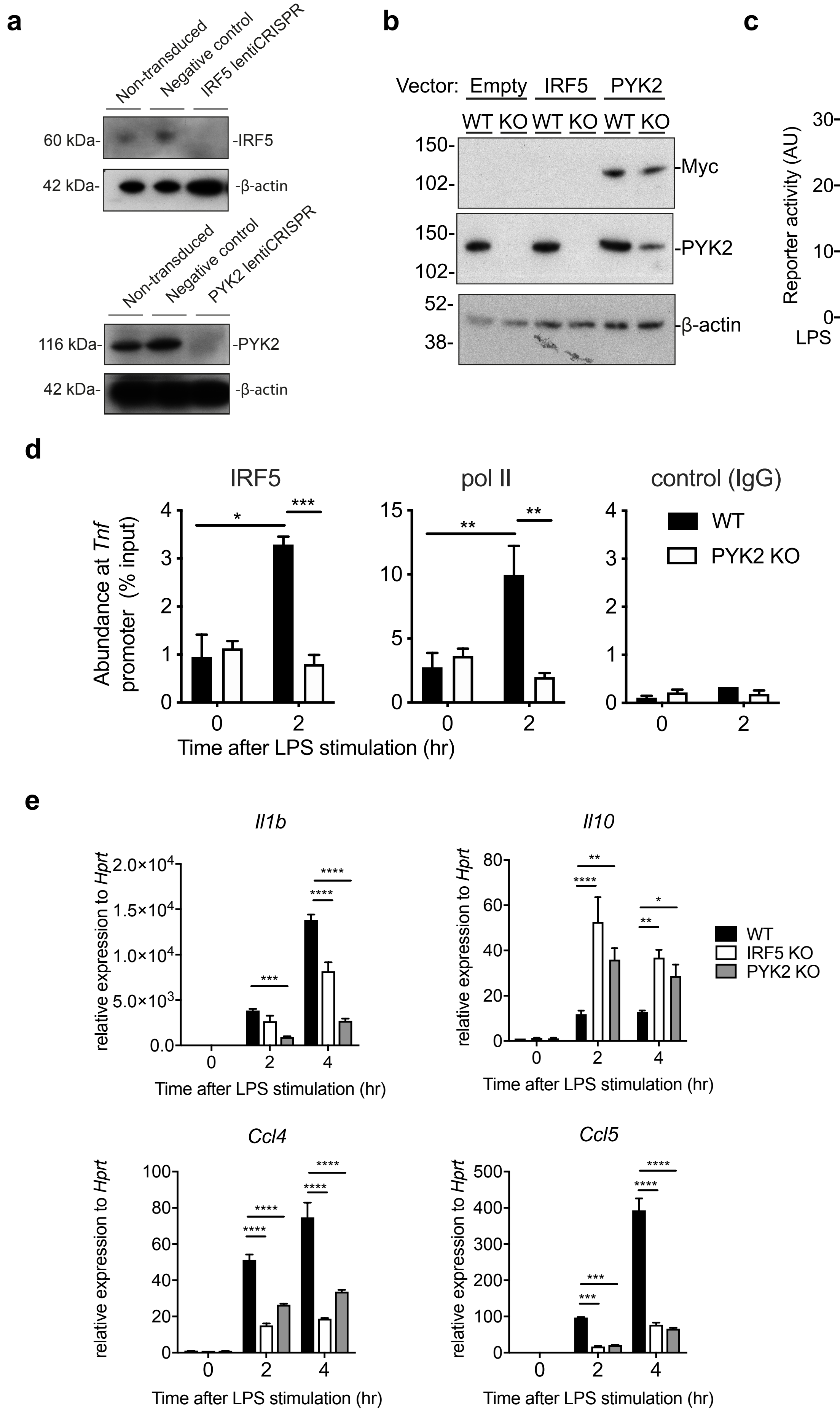

### Supplementary Fig. 3

#### Supplementary Figure S3

**a**

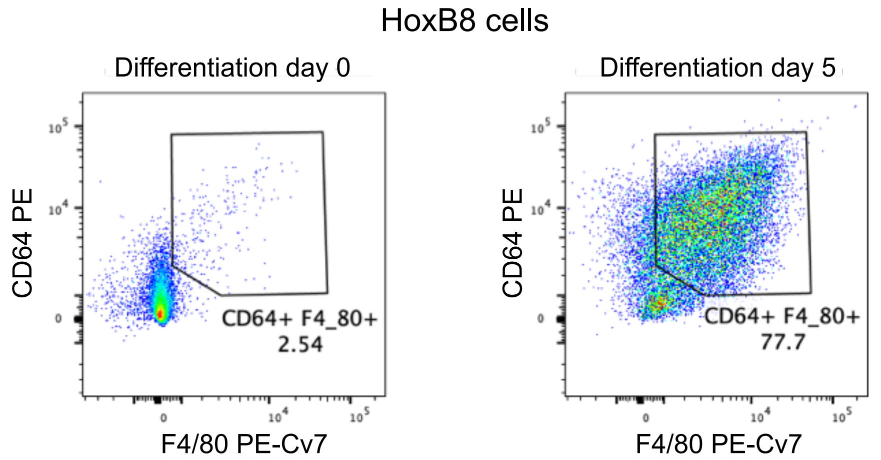


**b**

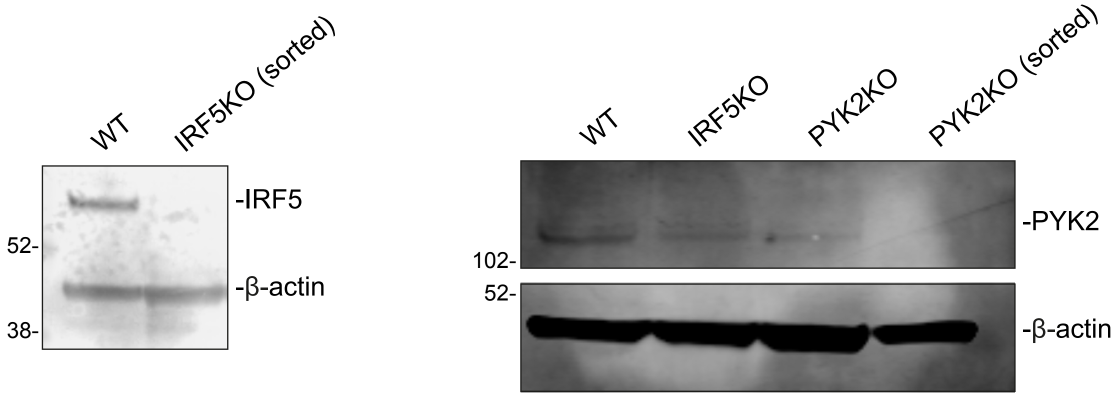


**c**

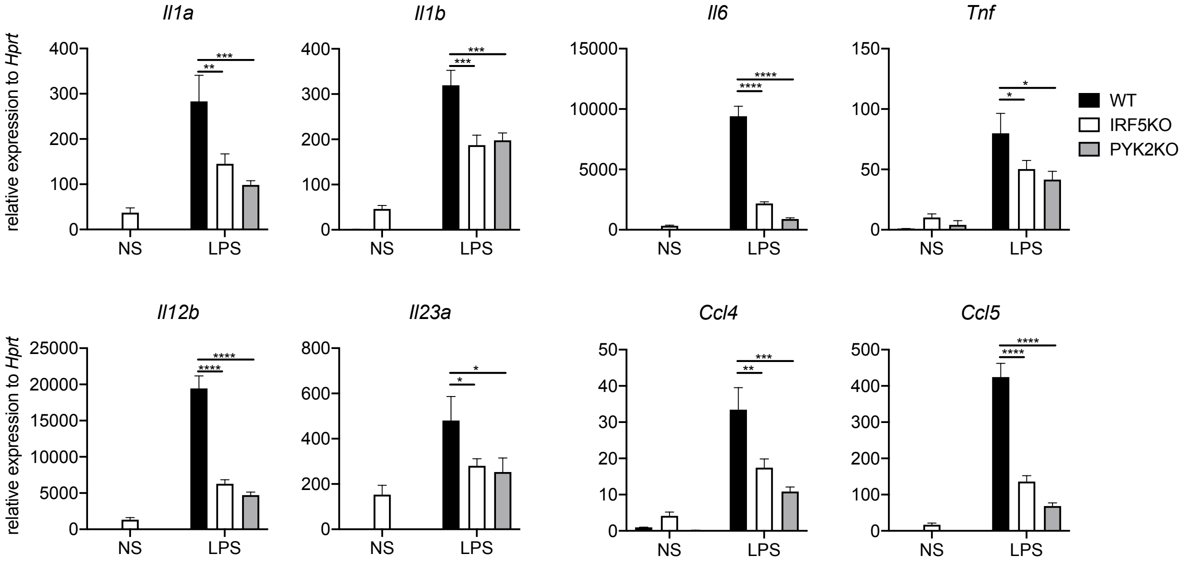

### Supplementary Fig. 4

#### Supplementary Figure S4

a

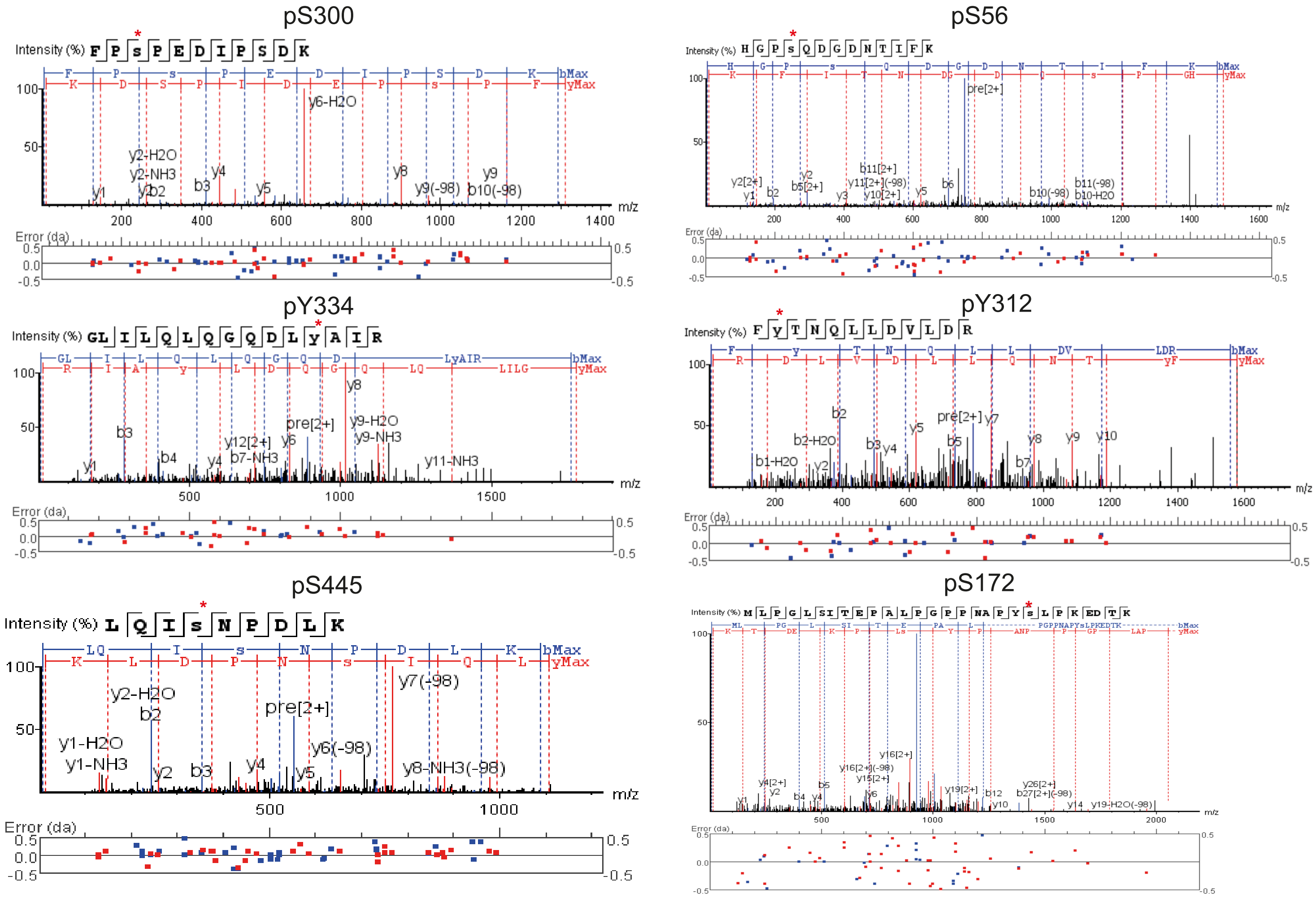


b

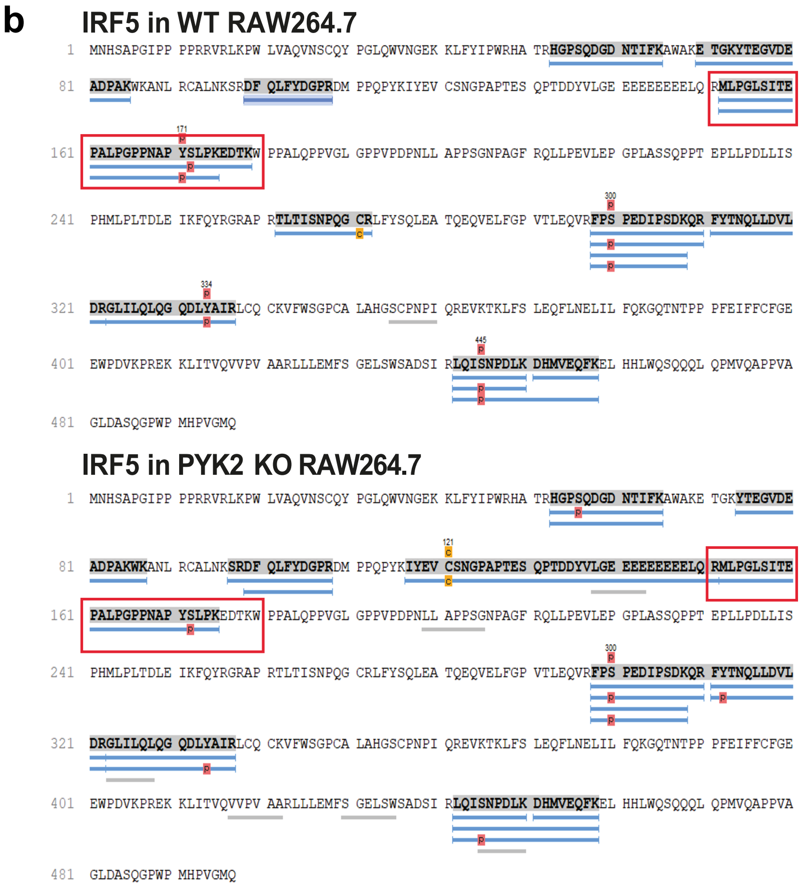


c

| Mouse IRF5 site | Human IRF5-v2 site |
|---|---|
| Y104 | Y104 |
| Y171 | Y172 |
| S172 | S173 |
| Y312 | Y329 |
| Y334 | Y351 |

### Supplementary Fig. 5

##### Supplementary Figure S5

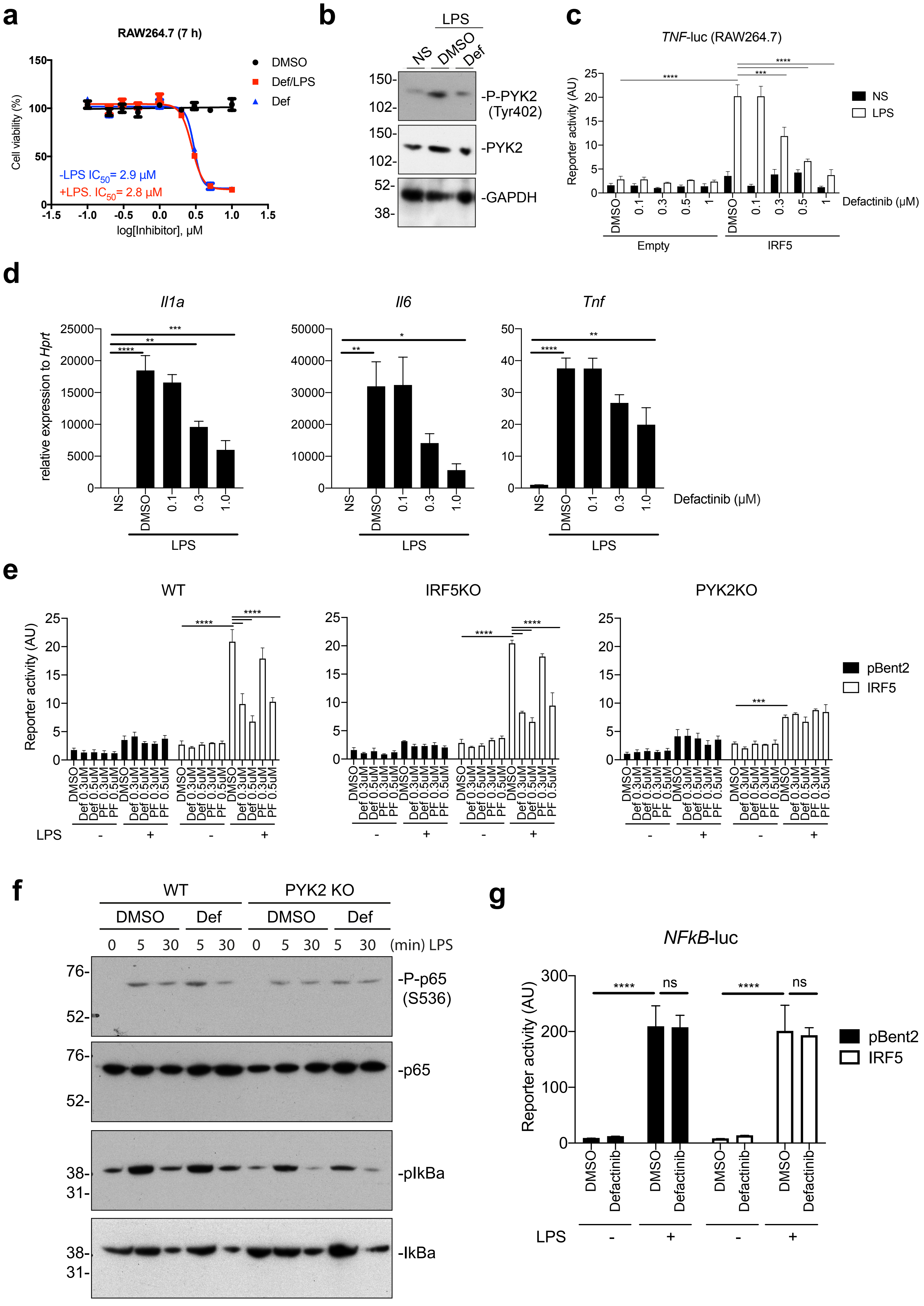

### Supplementary Fig. 6

#### Supplementary Figure S6

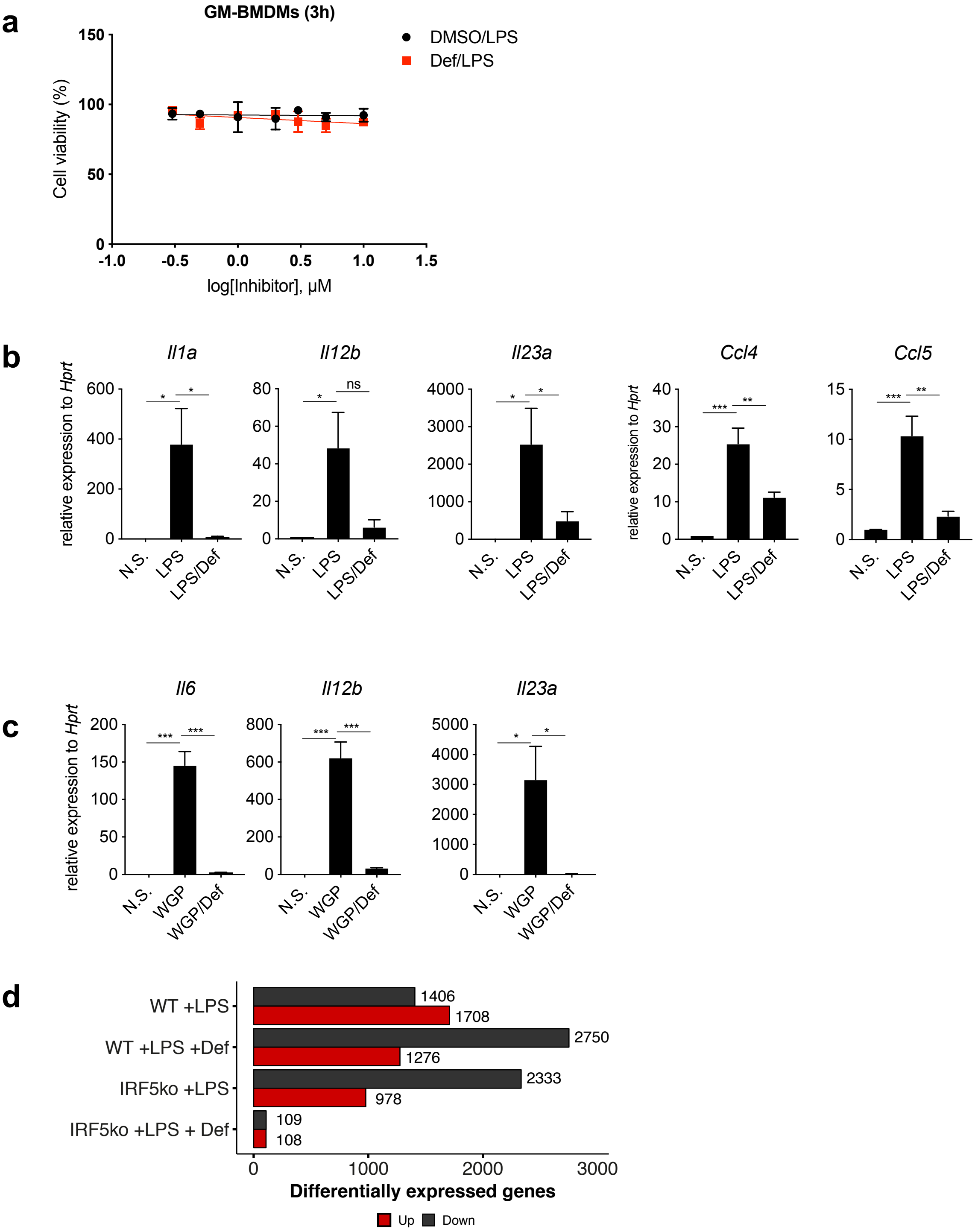

### Supplementary Fig. 7

#### Supplementary Figure S7

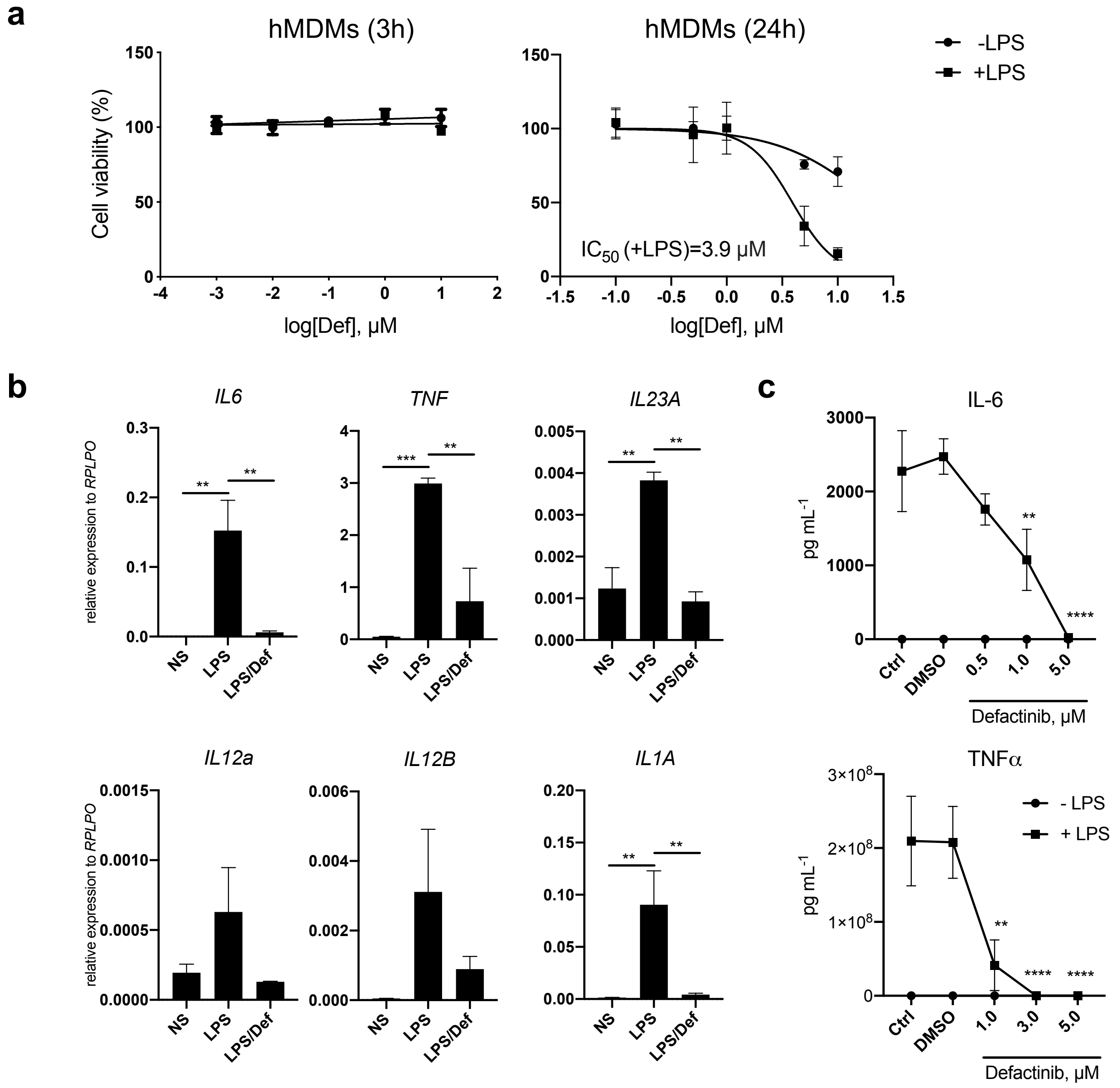

### Supplementary Fig. 8

#### Supplementary Figure S8

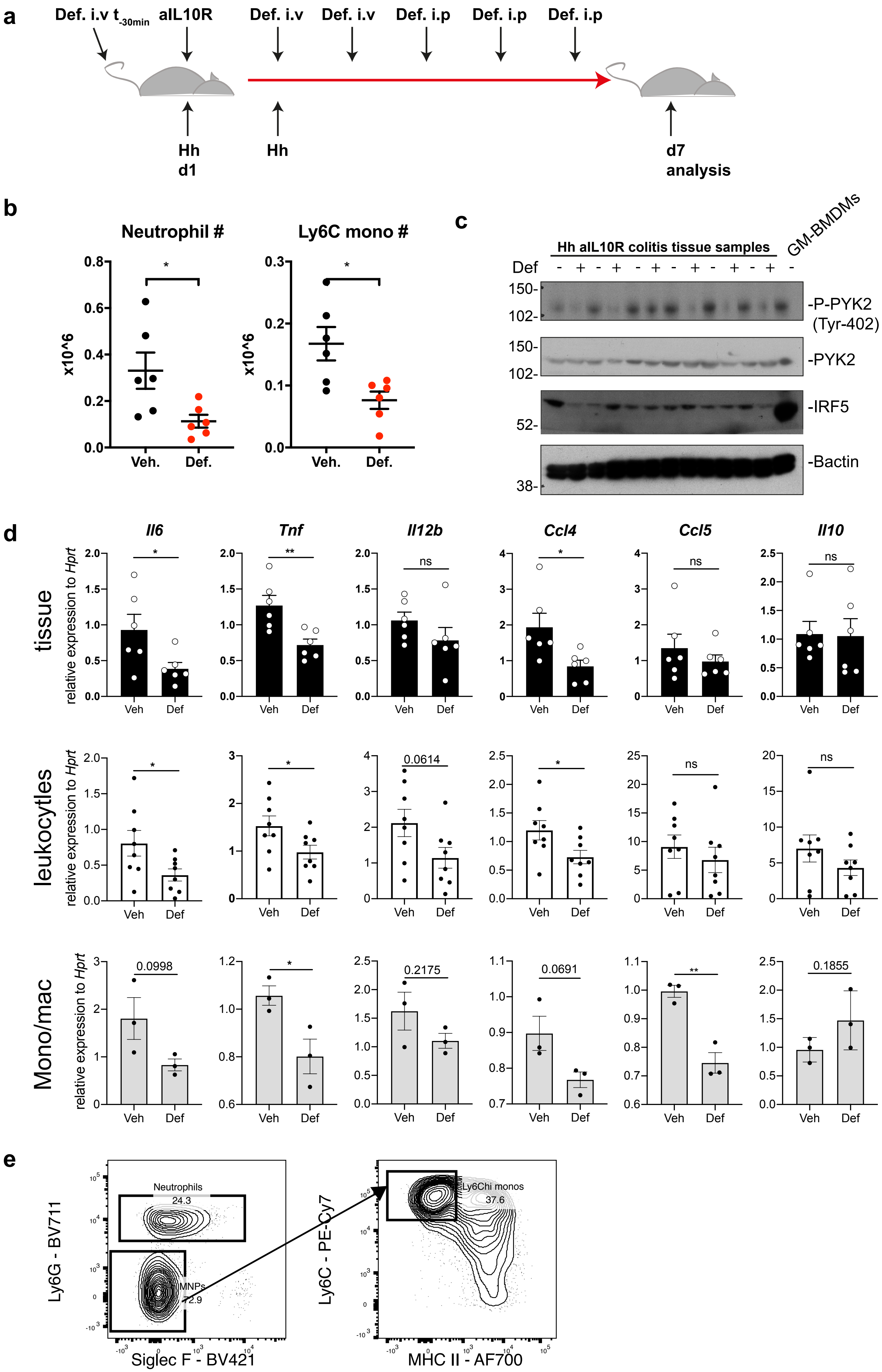

### Supplementary Fig. 9

**Supplementary Figure S9**

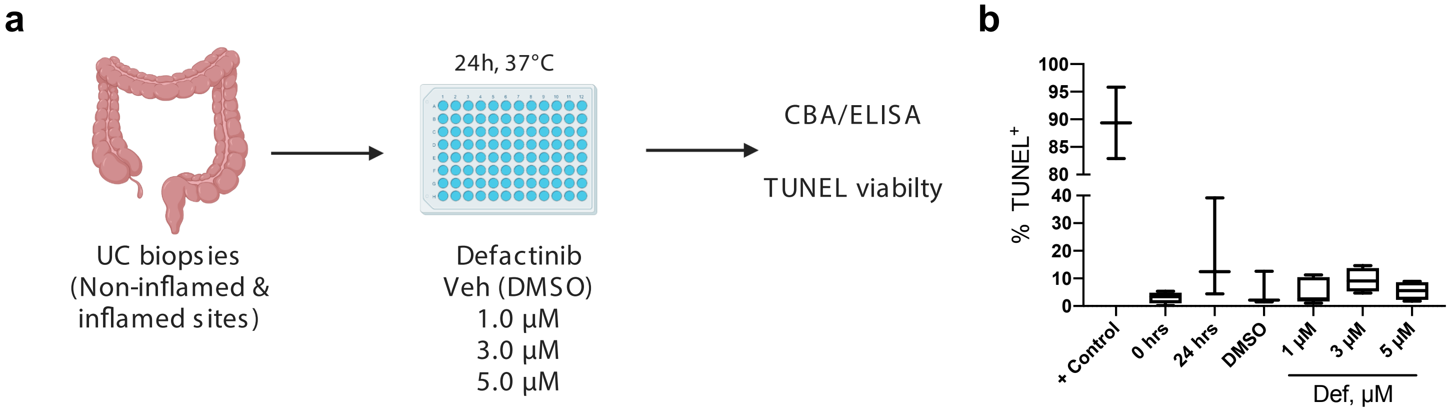

### Supplementary Fig. 10

#### Supplementary Figure S10

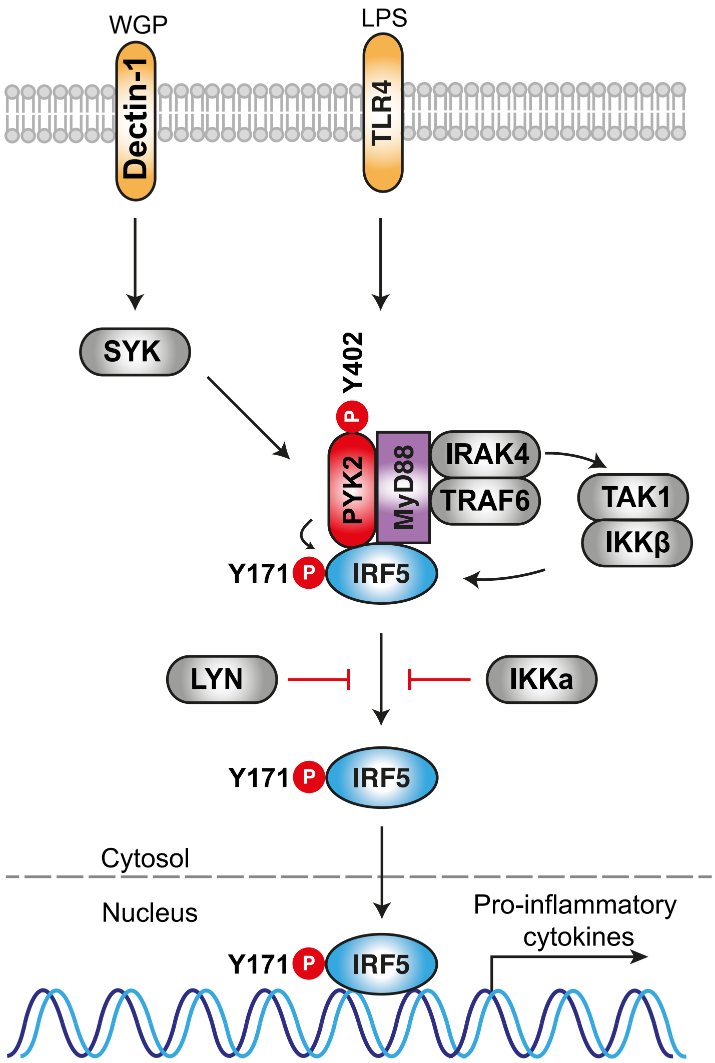
